# ΔN-Tp63 mediates Wnt/β-catenin-induced inhibition of differentiation in basal stem cells of mucociliary epithelia

**DOI:** 10.1101/682534

**Authors:** Maximilian Haas, José Luis Gómez Vázquez, Dingyuan Iris Sun, Hong Thi Tran, Magdalena Brislinger, Alexia Tasca, Orr Shomroni, Kris Vleminckx, Peter Walentek

## Abstract

Mucociliary epithelia provide a first line of defense against pathogens in the airways and the epidermis of vertebrate larvae. Impaired regeneration and remodeling of mucociliary epithelia are associated with dysregulated Wnt/β-catenin signaling in chronic airway diseases, but underlying mechanisms remain elusive and studies of Wnt signaling in mucociliary cells yield seemingly contradicting results. Employing the *Xenopus* mucociliary epidermis, the mouse airway, and human airway basal stem cell cultures, we characterize the evolutionarily conserved roles of Wnt/β-catenin signaling in mucociliary cells in vertebrates. Wnt signaling is required in multiciliated cells for cilia formation during differentiation stages, but in Basal cells, Wnt signaling prevents specification and differentiation of epithelial cell types by activating *ΔN-TP63* expression. We demonstrate that ΔN-TP63 is a master transcription factor in Basal cells, which is necessary and sufficient to mediate the Wnt-induced inhibition of differentiation and is required to retain basal stem cells during development. Chronic stimulation of Wnt signaling leads to mucociliary remodeling and Basal cell hyperplasia, but this is reversible *in vivo* and *in vitro*, suggesting Wnt inhibition as an option in the treatment of chronic lung diseases. Our work sheds light into the evolutionarily conserved regulation of stem cells and differentiation, resolves Wnt functions in mucociliary epithelia, and provides crucial insights into mucociliary development, regeneration and disease mechanisms.

## Introduction

Mucociliary epithelia line the conducting airways of most vertebrates as well as the epidermis of many vertebrate and invertebrate larvae (Walentek and Quigley, 2017). They are composed of multiple secretory cell types, including Goblet and outer cells, which release mucus, along with Ionocytes, Club cells and Small Secretory cells (SSCs), which release ions and small molecules into the extracellular space; in addition, multiciliated cells (MCCs) transport fluid along epithelia by coordinated ciliary motion, and Basal cells (BCs) reside underneath the epithelia and serve as tissue-specific stem cells (cf. graphical abstract) (Hogan et al., 2014; Rock et al., 2010). Mucociliary epithelia provide a first line of defense against pathogens by mucociliary clearance, which relies on the correct numbers and function of MCCs and secretory cells (Mall, 2008). Aberrations of cell type composition and BC behavior are observed in chronic lung diseases, e.g. chronic obstructive pulmonary disease (COPD), leading to impaired clearance and airway infections (Hogan et al., 2014; Tilley et al., 2014). While chronic lung diseases are among the most common causes of death worldwide, their pathogenic mechanisms are poorly understood and treatment options are very limited.

The plethora of diverse cell signaling functions is contrasted with a small number of pathways that are employed reiteratively to induce context-dependent responses. This complicates the interpretation of results from experimental manipulations of cell signaling in any given process or tissue. Wnt/β-catenin signaling regulates gene expression and plays a role in virtually all cells and tissues (Clevers, 2006). The pathway is activated by extracellular binding of Wnt ligands to Frizzled receptors and LRP5/6 co-receptors, which then recruit components of the β-catenin destruction complex including the kinase GSK3β to the membrane, where they are inhibited (Niehrs, 2012). β-catenin is then stabilized and enters the nucleus where it acts as transcriptional co-regulator through binding to TCF/LEF transcription factors.

Wnt/β-catenin signaling functions in mucociliary epithelia, but results from manipulations often appear contradictory as to the exact roles Wnt signaling plays in different cell types and processes. Wnt/β-catenin was suggested to promote MCC specification and expression of *FOXJ1*, a key transcription factor in motile cilia formation (Hou et al., 2019; Huang and Niehrs, 2014; Malleske et al., 2018; Stubbs et al., 2008; Walentek et al., 2012, 2015). In contrast, Wnt/β-catenin activation can also lead to loss of MCCs or Goblet secretory cell hyperplasia (Hashimoto et al., 2012; Mucenski et al., 2005; Reynolds et al., 2008; Schmid et al., 2017). Complicating matters further, additional effects for Wnt were proposed in submucosal glands, during regeneration and in regulating proliferation (Driskell, 2004; Hogan et al., 2014; Pongracz and Stockley, 2006). Dysregulation of Wnt signaling is commonly observed in chronic lung diseases such as COPD (Baarsma and Königshoff, 2017; Pongracz and Stockley, 2006). Thus, fundamental knowledge on the precise roles of Wnt/β-catenin in mucociliary cells is crucial to understand disease mechanisms and can provide entry points to develop treatments for patients.

We investigated the roles of Wnt/β-catenin in vertebrate mucociliary epithelia using the embryonic *Xenopus* epidermis, the mouse airway and human airway basal cell culture as models. Employing a combination of signaling reporter studies with single cell resolution, manipulations of the Wnt pathway during various phases of development and regeneration, and in epistasis experiments with downstream factors, we characterize the roles and effects of signaling on mucociliary cell types. Our data confirm a role of Wnt/β-catenin signaling in MCC differentiation, but also show its importance in the regulation of BCs. Collectively, we propose that high levels of Wnt/β-catenin signaling block differentiation of BCs into epithelial cell types by activating *ΔN-TP63* expression, which is necessary and sufficient to mediate this effect and to retain stem cells. Importantly, this inhibition of differentiation is reversible *in vivo* and *in vitro* suggesting local Wnt/β-catenin signaling manipulations to be further explored in the context of chronic lung diseases associated with airway epithelial remodeling.

## Results

### Wnt/β-catenin functions in MCCs and BCs

Wnt/β-catenin signaling was implicated in the specification and differentiation of secretory cells and MCCs in the mammalian airway as well as the *Xenopus* mucociliary epidermis, which serves as valuable model to investigate the principles of regulation and function of vertebrate mucociliary epithelia (Huang and Niehrs, 2014; Mucenski et al., 2005; Walentek et al., 2015). To clarify the roles of Wnt/β-catenin signaling in mucociliary cell types, we analyzed signaling activity using transgenic reporter lines expressing GFP upon Wnt/β-catenin activation in *Xenopus* and the mouse (Borday et al., 2018; Ferrer-Vaquer et al., 2010). Wnt signaling activity was assessed throughout development of the *Xenopus* epidermis and in the mouse conducting airways until mucociliary epithelia were fully mature **(Figure 1A,B and Supplemental Figures S1A and S2A)**. While the epidermis and the airways are derived from different germ layers and formed at different stages relative to organismal development (Walentek and Quigley, 2017; Warburton et al., 2010), our analysis revealed striking similarities in Wnt/β-catenin activity in both tissues. Initially, signaling activity was observed in cells throughout the epithelia, without particular compartmentalization. With progressive development, Wnt activity was restricted to the sensorial layer of the *Xenopus* epidermis **(Figure 1A)** and the basal compartment of the pseudostratified airway epithelium **(Figure 1B)**. In both systems, the location of Wnt signaling-positive cells coincided with the known location of the respective progenitor cell population that gives rise to MCCs and secretory cells, which then intercalate into the epithelium during differentiation (Deblandre et al., 1999; Rock et al., 2009; Stubbs et al., 2006). In *Xenopus*, we also observed GFP-positive cells in the epithelial cell layer during stages of intercalation (st. 25) **(Figure 1A**; arrowheads). En-face imaging in combination with immunostaining for cell-type markers revealed increased Wnt activity in intercalating MCCs and Ionocytes at stage 25 **(Supplemental Figure S1B)**. In the mature mucociliary epidermis, Wnt activity was then restricted to MCCs **(Figure 1C)**. We also detected elevated Wnt activity in differentiating MCCs of the mouse airway, although reporter activity was lower in MCCs as compared to Wnt-positive cells residing at the base of the epithelium **(Figure 1D and Supplemental Figure S2B)**. We generated mouse tracheal epithelial cell (MTEC) cultures from Wnt-reporter animals and monitored Wnt activity in the air-liquid interface (ALI) *in vitro* regeneration model at days 1, 4, 7, 14 and 21 (Vladar and Brody, 2013). Wnt signaling activity was detected throughout all stages of regeneration, with MCCs showing elevated signaling levels as well as reporter-positive cells residing basally, but no Wnt activity was detected in Club cells **(Supplemental Figure S2C-E)**.

**Figure 1:**
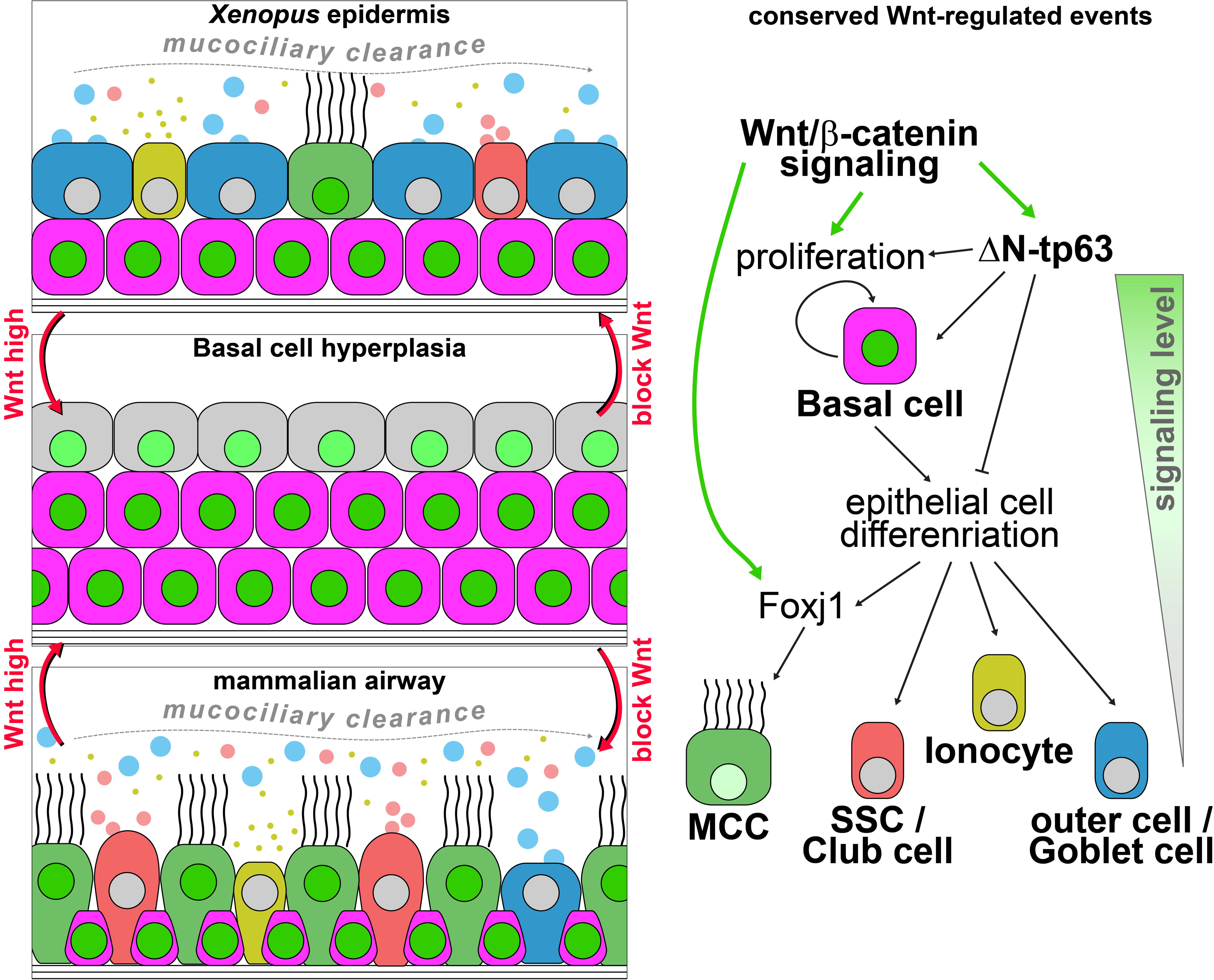
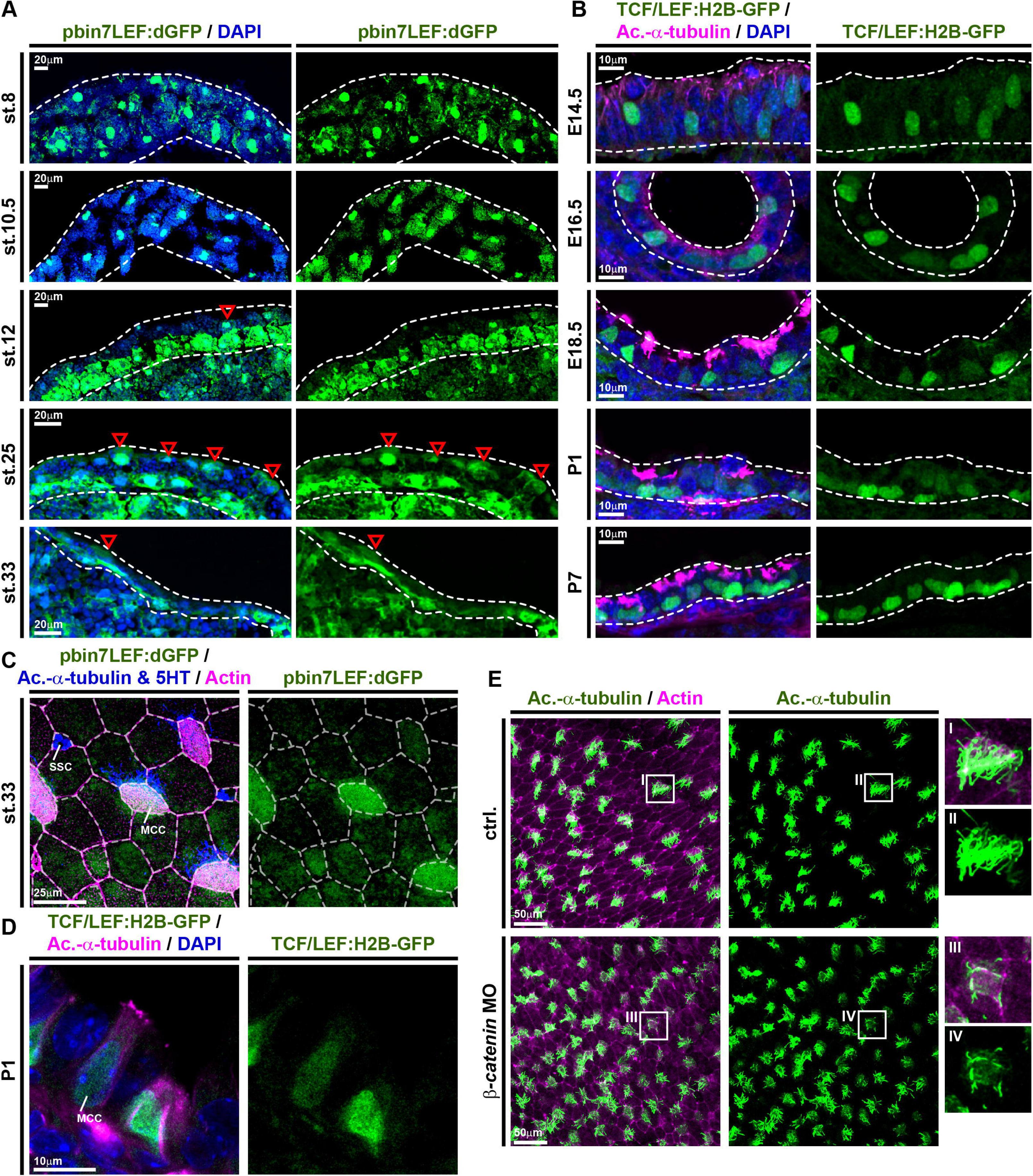
Wnt/β-catenin signaling is active in MCCs and basal progenitors. **(A)** Analysis of Wnt/β-catenin activity in the *X. laevis* mucociliary epidermis using the pbin7LEF:dGFP reporter line (green). Nuclei are stained by DAPI (blue). Red arrowheads indicate GFP-positive cells in the outer epithelial layer. Dashed lines outline the epidermal layers. Embryonic stages (st. 8-33) are indicated. **(B)** Analysis of Wnt/β-catenin activity in the mouse developing airway mucociliary epithelium using the TCF/LEF:H2B-GFP reporter line (green). Nuclei are stained by DAPI (blue) and MCCs are marked by Acetylated-α-tubulin (Ac.-α-tubulin, magenta) staining. Dashed lines outline the epithelium. Embryonic (E14.5-18.5) / post-natal (P1-7) stages are indicated. **(C)** En-face imaging of the mature *Xenopus* epidermis at st. 33 shows elevated signaling levels (green) in MCCs (Ac.-α-tubulin, blue). SSCs (5HT, blue). Cell membranes are visualized by Actin staining (magenta). **(D)** Immunostaining for Ac.-α-tubulin (magenta) and nuclei (DAPI, blue) shows high levels of Wnt signaling (green) in cells with BC morphology and intermediate signaling levels in differentiating MCCs. **(E)** Morpholino-oligonucleotide (MO) knockdown of *β-catenin* in *Xenopus* increases MCC numbers (Ac.-α-tubulin, green), but MCCs present fewer and shorter cilia than controls (ctrl.). Actin staining (magenta). Insets indicate locations of magnified areas I-IV. **(Related to Supplementary Figures S1 and S2)**

These data suggested a role for Wnt/β-catenin in basal progenitor cells as well as in MCCs. To test this, we knocked down β-catenin using morpholino oligonucleotide (MO) injections targeting the *Xenopus* epidermis, and analyzed epidermal morphology as well as MCCs **(Figure 1E)**. We observed increased numbers of MCCs in *β-catenin* morphants (*β-catenin* MO), but these MCCs presented reduced numbers of cilia **(Figure 1E, Supplemental Figure 1C)**. These data resembled experiments using overexpression of the LRP6-inhibitor *dickkopf 1* (*dkk1*) in *Xenopus* (Walentek et al., 2015). Reduced ciliation rate in β-catenin-deficient MCCs was also compatible with data demonstrating that β-catenin is a transcriptional co-regulator of *foxj1*, which is required for motile ciliogenesis in all vertebrate MCCs (Caron et al., 2012; Gomperts, 2004; Stubbs et al., 2008; Walentek et al., 2012). Nevertheless, the question arose as to why reduced β-catenin levels increased the overall number of MCCs in the epithelium. As the basal precursor cell compartment was the site of highest Wnt signaling reporter-activity in both *Xenopus* and mice, we wondered if loss of β-catenin would affect BCs and lead to increased MCC specification. We injected *β-catenin* MO targeting exclusively the right side of embryos, and analyzed marker gene expression for MCCs and BCs at mid-neurula stages (st. 17), i.e. after cell fate specification. *In situ* hybridization (ISH) for *foxj1* (MCCs) and *ΔN-tp63* (sensorial layer BCs; (Cibois et al., 2015; Lu et al., 2001)) revealed an increase in *foxj1*-positive cells and reduced *ΔN-tp63* expression on the injected side of the embryos **(Supplemental Figure 1D,E)**. This implicated an increase in MCC specification at the expense of basal progenitors upon Wnt inhibition. Collectively, our experiments identified MCCs and BCs as sites of elevated signaling activity during mucociliary development, and a requirement for controlled Wnt/β-catenin signaling in MCCs and BCs to generate a normal mucociliary epithelium.

### ΔN-Tp63 is necessary and sufficient to block differentiation in response to Wnt/β-catenin

Airway BCs are tissue-specific stem cells and required for maintenance and regeneration of all mucociliary cell types (Rock et al., 2010; Zuo et al., 2015). ΔN-TP63-α is the dominantly expressed isoform of the transcription factor TP63 in airway BCs and a commonly used marker for BCs in various epithelia (Arason et al., 2014; Soares and Zhou, 2018; Warner et al., 2013). Expression of *ΔN-TP63* isoforms is regulated by an evolutionarily conserved alternative promotor (P2) initiating transcription at alternative exon 3 (Ruptier et al., 2011). In *Xenopus*, only *ΔN-tp63* isoforms are expressed during development, and no full-length isoform is annotated in the *X. laevis* or *X. tropicalis* genomes to date, indicating potential loss of this isoform in the frog. Nevertheless, in *Xenopus* and in mammals, chromatin immunoprecipitation and DNA-sequencing (ChIP-seq) has detected multiple TCF/LEF binding sites in P2, suggesting direct Wnt/β-catenin regulation (Kjolby and Harland, 2017; Ruptier et al., 2011). Since *ΔN-TP63* is associated with the regulation of differentiation and given our observation that loss of β-catenin lead to decreased *ΔN-tp63* expression, we tested if *ΔN-tp63* was Wnt-regulated in the mucociliary epidermis. Ectopic activation of Wnt/β-catenin signaling was achieved by application of the GSK3β-inhibitor 6-Bromoindirubin-3’-oxime (BIO) to the medium starting at st. 8 of *Xenopus* development. Efficient activation of the Wnt pathway in the epidermis was confirmed using the Wnt-reporter line **(Figure 2A)**. First, we analyzed the effects of BIO treatment on epidermal *ΔN-tp63* expression by ISH. Specimens treated with BIO displayed increased levels of *ΔN-tp63* expression and a thickening of the sensorial layer **(Figure 2B and Supplemental Figure S3A)**. Next, we treated embryos with BIO from st. 8 until st. 30, when MCCs and Ionocytes have fully developed, and analyzed cell type composition by immunofluorescent staining (Walentek, 2018). Treatment with BIO significantly reduced the numbers of all intercalating cell types in a dose-dependent manner **(Figure 2C and Supplemental Figure S3B,C)**. Lack of mature MCCs and Ionocytes could be a result of inhibited cell fate specification or defective differentiation and intercalation into the epithelium. Therefore, we also tested the effects of BIO treatment on the expression of early cell type-markers associated with successful cell fate specification by ISH at st. 17, i.e. before intercalation (Walentek and Quigley, 2017). We observed a loss or strong reduction in cell type-marker expression for MCCs (*foxj1*), Ionocytes (*foxi1*; (Quigley et al., 2011)) and SSCs (*foxa1*; (Dubaissi et al., 2014)), indicating a failure in cell fate specification after BIO application **(Figure 2D and Supplemental Figure S3D,E)**. These data suggested that increased Wnt/β-catenin lead to upregulation of *ΔN-tp63* and expansion of the BC pool, while inhibiting specification of epidermal cell types. To directly test if ΔN-tp63 was necessary for the block of specification in response to Wnt overactivation, we injected embryos with a *ΔN-tp63* MO at four-cell stage and treated the morphants either with vehicle or BIO, starting at st. 8. Cell type quantification at st. 30 and ISH marker analysis at st. 17 both showed a partial rescue of cell fate specification and morphogenesis in *ΔN-tp63* MO embryos treated with BIO, and an increased specification of MCCs and Ionocytes in *ΔN-tp63* MO morphants without BIO application **(Figure 2C,D and Supplemental Figure S3B-E)**. Together, these results indicated that *ΔN-tp63* activation was necessary to block differentiation upon BIO treatment. Next, we wondered if ΔN-tp63 alone was sufficient to inhibit differentiation in the absence of increased Wnt signaling. Therefore, we generated GFP-tagged and untagged ΔN-tp63 constructs. Overexpression of *gfp-ΔN-tp63* in the epidermis and immunofluorescent staining confirmed successful production of the protein and its nuclear localization **(Figure2 E)**. Furthermore, injections of *gfp-ΔN-tp63* or *ΔN-tp63* reduced MCC numbers and expression of early cell type markers for MCCs, Ionocytes and SSCs **(Figure2 E,F and Supplemental Figure S4A-C)**, thereby providing evidence for sufficiency. Additionally, we investigated if ΔN-tp63 only inhibits cell fate specification from BCs or if its activity in cells after specification could inhibit differentiation as well. For that, we generated a hormone-inducible version of GFP-ΔN-tp63 (GFP-ΔN-tp63-GR; (Kolm and Sive, 1995)), injected embryos at four-cell stage, and added Dexamethasone (Dex) to the medium at various stages of development. Activation of the construct and subsequent nuclear localization was confirmed by confocal microscopy **(Supplemental Figure S4D)**. Dex addition at st. 9 suppressed MCC formation as observed with the non-inducible construct, whereas application of vehicle at st. 9 or Dex activation after MCC specification at st. 24 did not result in reduced MCC numbers at st. 30 **(Supplemental Figure S4E,F)**. High-magnification imaging further confirmed presence of GFP-ΔN-tp63-GR in the nuclei of fully differentiated MCCs and Ionocytes in specimens activated at st. 24 **(Supplementary Figure S4G)**. These results indicated that the inhibitory effect of ΔN-tp63 on epithelial cell specification was restricted to basal progenitors. Finally, we investigated the degree of evolutionary conservation of the observed effects in human airway basal stem cells. Ectopic activation of canonical Wnt signaling in ALI cultures derived from immortalized human airway BCs (BCi-NS1.1 cells, BCIs (Walters et al., 2013)) was induced by application of human recombinant R-spondin 2 (RSPO2) protein to the medium after initial epithelialization of cultures was completed at ALI day 7. We then analyzed the effects of RSPO2 on airway mucociliary regeneration by immunofluorescent staining and quantitative RT-PCR (qPCR). In BCIs, RSPO2 application inhibited differentiation of MCCs and Club secretory cells **(Figure 3A,C,D)**. At the same time, we observed an increase in *ΔN-TP63* expression after RSPO2 treatment as well as elevated levels for *KRT5* (*Keratin 5*), an additional marker for BCs in airway epithelia **(Figure 3C-E)** (Zuo et al., 2015). Orthogonal optical sections of confocal images from BCIs stained for ΔN-TP63 or KRT5 further revealed an increase in epithelial thickness and epithelial KRT5-positive cells after RSPO2 treatment **(Figure 3E,F and Supplemental Figure S5A,B)**, similar to our observations in *Xenopus* and reminiscent of phenotypes in COPD patients (Rock et al., 2010), potentially indicating early BC hyperplasia. This interpretation of results was further supported by quantification of Ki67-positive proliferative cells, which number increased upon RSPO2 treatment, while the total number of epithelial cells remained low, likely due to inhibited specification and intercalation of MCCs and Club cells into the epithelium **(Supplemental Figure S5C-E)**. To address if Goblet cell hyperplasia occurred after Wnt/β-catenin gain-of-function in addition to effects on MCC and Club cell specification, we also analyzed Mucin expression. qPCR on BCIs treated with RSPO2 revealed reduced *MUC5A/C* expression, while the expression of *MUC5B* was elevated **(Supplemental Figure S5F,G)**. Nevertheless, immunofluorescent staining did not detect an increase in epithelial cells staining for MUC5B **(Supplemental Figure S5H)**. In summary, our data revealed that ΔN-TP63 was necessary and sufficient to inhibit differentiation of mucociliary epithelial cell types from BCs in response to canonical Wnt activation, without the need for Goblet cell hyperplasia. Furthermore, our results suggested that prolonged overactivation of Wnt signaling could lead to BC hyperplasia and long-term remodeling of the airway epithelium.

**Figure 2:**
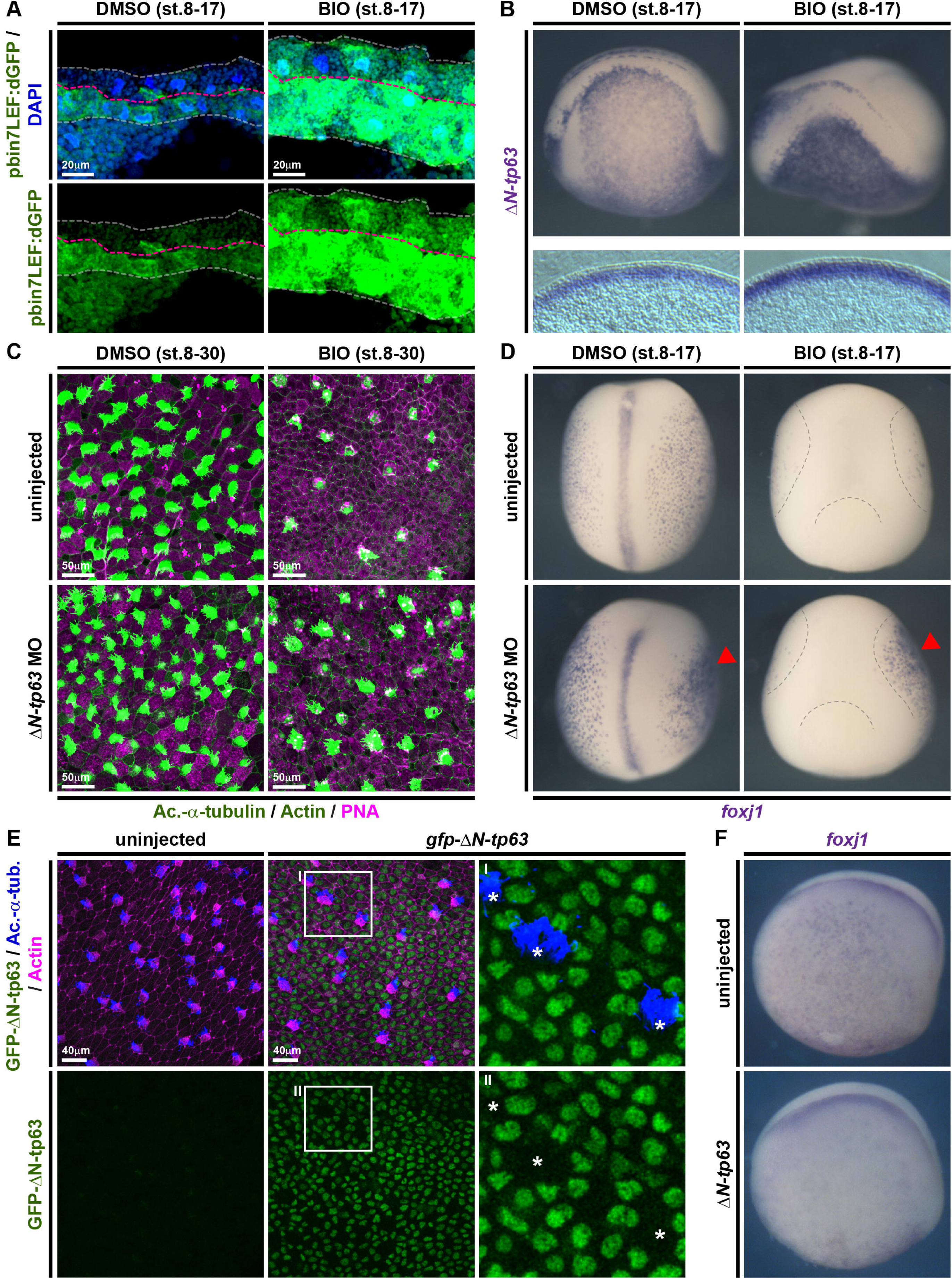
ΔN-tp63 mediates Wnt/β-catenin-induced inhibition of cell fate specification in *Xenopus*. **(A-D)** BIO treatments from st. 8-17 or st. 8-30. DMSO was used as vehicle control. **(A)** Confocal imaging shows Wnt/β-catenin signaling-activation (green) and thickening of the epidermis in BIO treated embryos. Nuclei (DAPI, blue). DMSO (N=2), BIO (N=2). **(B)** *In situ hybridization* (ISH) shows increased intensity and thickness of the *ΔN-tp63* expression domain in BIO treated whole mounts (upper row) and transversal sections (bottom row). **(C)** BIO treatment reduces MCC (Ac.-α-tubulin, green), Ionocyte (no staining, black), and SSC (large vesicles, PNA staining, magenta) numbers in confocal micrographs. Actin staining (green). *ΔN-tp63* MO in controls leads to increased MCCs and Ionocytes, while *ΔN-tp63* MO in BIO treated embryos rescues MCC and Ionocyte formation. **(D)** ISH reveals reduced MCC numbers (*foxj*+ cells) after BIO treatment, while unilateral knockdown of *ΔN-tp63* in control treated embryos leads to more *foxj*+ cells and rescues *foxj*+ cell number in BIO treated embryos. DMSO (N=5), BIO (N=7), DMSO+*ΔN-tp63* MO (N=5), BIO+*ΔN-tp63* MO (N=4). Injected side is indicated by red arrowhead. Dashed lines indicate epidermal area in BIO treated embryos. **(E)** Overexpression of *gfp-ΔN-tp63* mRNA leads to nuclear localization of the protein (green) and reduced MCC (Ac.-α-tubulin, blue) numbers at st. 30. Actin staining (magenta). Differentiated MCCs in injected specimens show no nuclear GFP signal (asterisks), indicating that they were not targeted. Magnified areas I-II are indicated. **(F)** ISH for *foxj1* at st. 17 shows reduced MCCs in *ΔN-tp63* mRNA injected embryos. **(Related to Supplementary Figures S3 and S4)**

**Figure 3:**
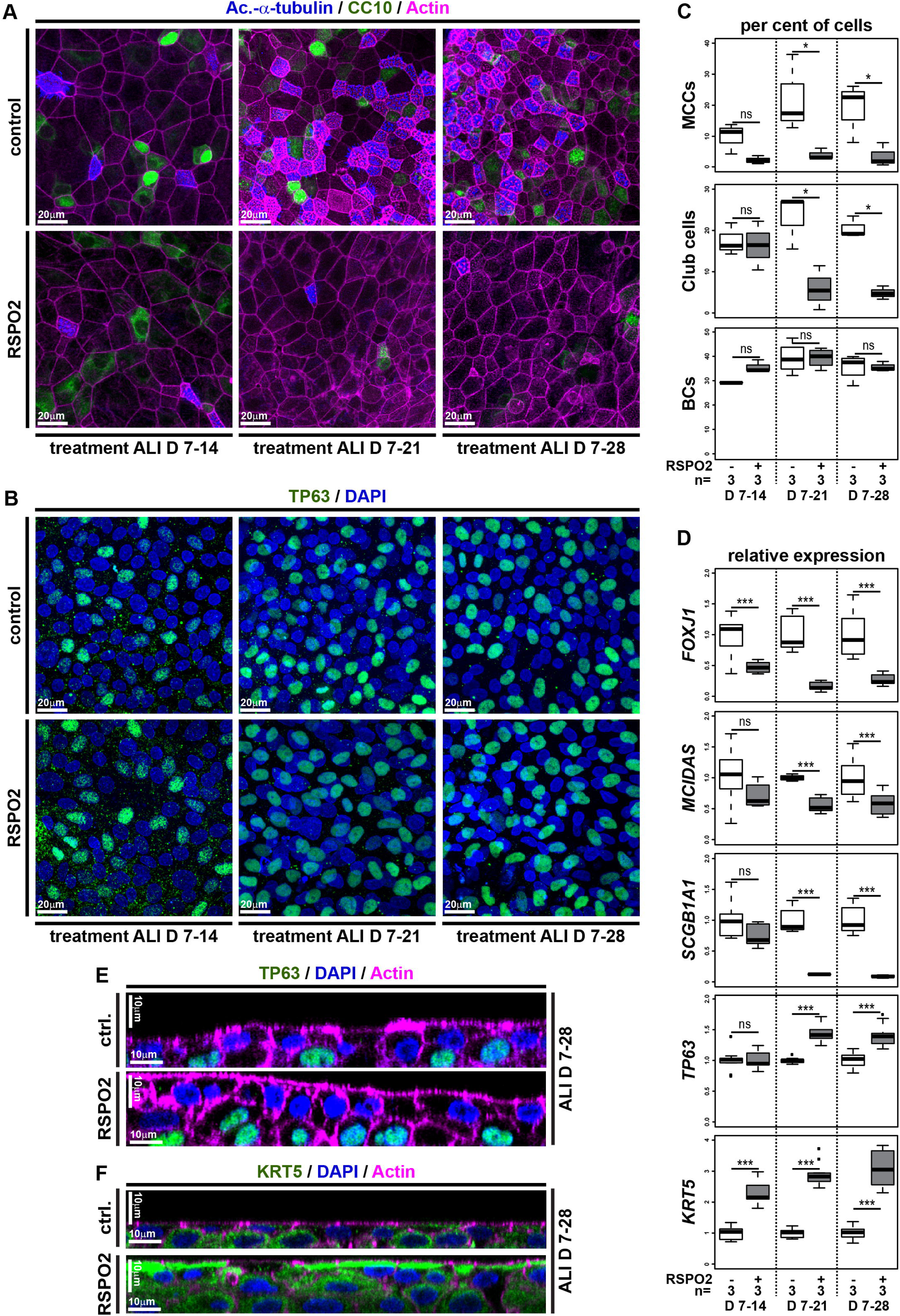
Wnt/β-catenin signaling inhibits differentiation and promotes stemness in human BCs. **(A-F)** Human immortalized BC (BCIs) kept in air-liquid interface (ALI) culture for up to 4 weeks. Human recombinant R-spondin2 (RSPO2) was used to activate Wnt/β-catenin signaling starting at ALI day 7 (D7). N=3 cultures per timepoint and treatment. **(A)** Confocal imaging of specimens stained for Ac.-α-tubulin (MCCs, blue), CC10 (Club cells, green) and Actin (cell membranes, magenta) show reduced MCC and Club cell numbers in RSPO2-treated cultures. **(B)** RSPO2 does not reduce the number of TP63+ (green) cells. Nuclei (DAPI, blue). **(C)** Quantification from (A,B). Mann Whitney test, not significant, ns = *P*>0.05; * = *P*≤0.05; ** = *P*≤0.01; *** = *P*≤0.001. **(D)** Quantitative RT-PCR (qPCR). Expression levels are depicted relative to stage controls. RSPO2 reduces expression of MCC (*FOXJ1*, *MCIDAS*) and Club cell (*SCGB1A1*) markers, but increases expression of BC markers (*TP63*, *KRT5*). Student’s t-test, not significant, ns = *P*>0.05; * = *P*≤0.05; ** = *P*≤0.01; *** = *P*≤0.001. **(E,F)** Optical orthogonal sections of confocal images. RSPO2-treated cultures display increased epithelial thickness and cells staining for BC markers TP63 (green, in E) and KRT5 (green, in F). **(Related to Supplemental Figure S5)**

### ΔN-Tp63 and Wnt signaling are required for maintenance of BCs and correct cell type composition in mucociliary epithelia

While our results argued for an important role for ΔN-TP63 in BCs of mucociliary epithelia, we found it astonishing that developmental loss of ΔN-TP63 in mammals (Daniely et al., 2004) and *Xenopus* (this study) still allowed for the formation of a mucociliary epithelium. We therefore tested how *ΔN-tp63* knockdown affected the mucociliary epidermis in more detail. In contrast to MCCs and Ionocytes, which intercalate early (st. 25) and are fully mature by st. 30, SSCs appear and mature slightly later (Walentek et al., 2014), resulting in approximately equal numbers of MCCs, Ionocytes and SSCs in the mucociliary epidermis by st. 34 **(Figure 4C)** (Walentek, 2018). Injections of *ΔN-tp63* MO and subsequent analysis of cell type-composition at st. 34 revealed a moderate increase in MCCs and Ionocytes, but a significant decrease in SSCs **(Figure 4A,C,D)**. These results suggested that premature release of BCs into differentiation could have reduced the availability of BCs during later stages of SSC specification. Therefore, we tested if SSCs are indeed specified after MCCs and Ionocytes or if their late appearance in the epithelium could be a consequence of prolonged differentiation or slower intercalation. For that, embryos were treated with BIO, starting at st. 11. This later stimulation of Wnt/β-catenin signaling resulted in almost normal MCC and Ionocyte numbers, but completely inhibited the appearance of SSCs **(Figure 4B-D)**. These data strongly indicated that SSCs were derived from the same BC progenitor pool during development as MCCs and Ionocytes, and that SSCs were specified later. To test if *Xenopus* BCs were lost by *ΔN-tp63* MO, we generated mucociliary organoids from *Xenopus* animal cap explants, providing pure mucociliary tissue for RNA-sequencing (RNA-seq) (Walentek and Quigley, 2017). Organoids were generated from control and *ΔN-tp63* morphant embryos and collected for total RNA extraction at early (st. 10) and late (st. 17) cell fate-specification stages as well as after specification was completed (st. 25). RNA-seq and differential expression analysis revealed significant differences in gene expression between control and *ΔN-tp63* morphant samples, and the most differentially expressed genes were detected at st. 25 (243 genes with P-adj<0.05; **Supplementary Figure S6A and Table 1**) (Love et al., 2014). Among significantly upregulated genes, we found MCC and Ionocyte genes including *mcidas*, *ccno*, *cdc20b*, *foxn4* and *foxi1*, and Go-term analysis indicated enrichment for “multi-ciliated epithelial cell differentiation” **(Supplemental Table 1 and Figure S6B)** (Mi et al., 2013; Walentek and Quigley, 2017). In contrast, Go-term analysis of significantly downregulated genes identified an enrichment for the terms “focal adhesion”, “actin cytoskeleton” and “extracellular matrix” **(Supplemental Figure S6B)**. These terms were also found to be enriched within the human airway BC transcriptome, suggesting loss of BCs (Hackett et al., 2011). Next, we compared the list of differentially expressed genes in *ΔN-tp63* morphants with the human airway BC transcriptome and identified 41 dysregulated *Xenopus* homologues, including multiple regulators of proliferation and of cell/extracellular matrix interactions **(Figure 4E)**. We subjected their and *ΔN-tp63*’s relative expression values (log2 fold-change relative to controls) to hierarchical clustering. In our analysis, we also included a subset of previously identified *Xenopus* core MCC and Ionocyte markers as well as known markers for SSCs and Goblet/outer-layer cells (Dubaissi et al., 2014; Hayes et al., 2007; Quigley and Kintner, 2017). The two clusters representing the most upregulated genes over developmental time in *ΔN-tp63* morphants contained key markers for MCCs (e.g. *mcidas*, *ccno*, *cdc20b*, *foxj1*, *myb*) and Ionocytes (e.g. *foxi1*, *atp6* subunits, *ca12*, *ubp1*) **(Figure 4E)**. In contrast, the cluster representing the most downregulated genes over developmental time contained exclusively BC markers (e.g. *ΔN-tp63*, *itga3/6*, *itgb1*, *lamb1*, *cav2*), and the key transcription factor for SSC specification *foxa1* **(Figure 4E)**. Together, these data not only revealed a high degree of functional and transcriptional homology between human and *Xenopus* mucociliary BCs, but also demonstrated that ΔN-tp63 was necessary for the maintenance of BCs during development, which was in turn required to generate a normal cell type-composition in the mucociliary epidermis. Furthermore, these data were in line with previous work, demonstrating that loss of ΔN-TP63 impaired regeneration and induced senescence in human ALI cultures (Arason et al., 2014). Therefore, we wondered if elevated Wnt/β-catenin signaling in the airway basal compartment was required for the maintenance *ΔN-TP63* expression and BCs in human cells after epithelialization as well. To test this, we inhibited Wnt/β-catenin signaling by addition of human recombinant DKK1 protein (DKK1) to the medium of differentiating BCI ALI cultures starting at ALI day 7. Quantification of cell type markers and mRNA expression levels by immunofluorescence and qPCR showed a moderate increase in MCCs and Club cells and a relative decrease in BCs in DKK1-treated cultures, but not a long-term loss of BCs **(Figure 5B-F)**. Furthermore, proliferation and the total number of epithelial cells remained unchanged, and Mucin production was not inhibited after DKK1 treatment **(Supplemental Figure S6C-H)**. These data indicated that Wnt/β-catenin regulated the decision between BC identity and differentiation into epithelial cell types, but that it was dispensable in later phases of *in vitro* regeneration for maintenance of *ΔN-TP63* expression and BCs. In summary, our experiments demonstrated that ΔN-TP63 was a master transcription factor in BCs regulating the decision between differentiation and basal stem cell fate in vertebrate mucociliary epithelia, and that Wnt/β-catenin was required for maintaining *ΔN-TP63* expression and BCs during development, but not in later phases of regeneration.

**Figure 4:**
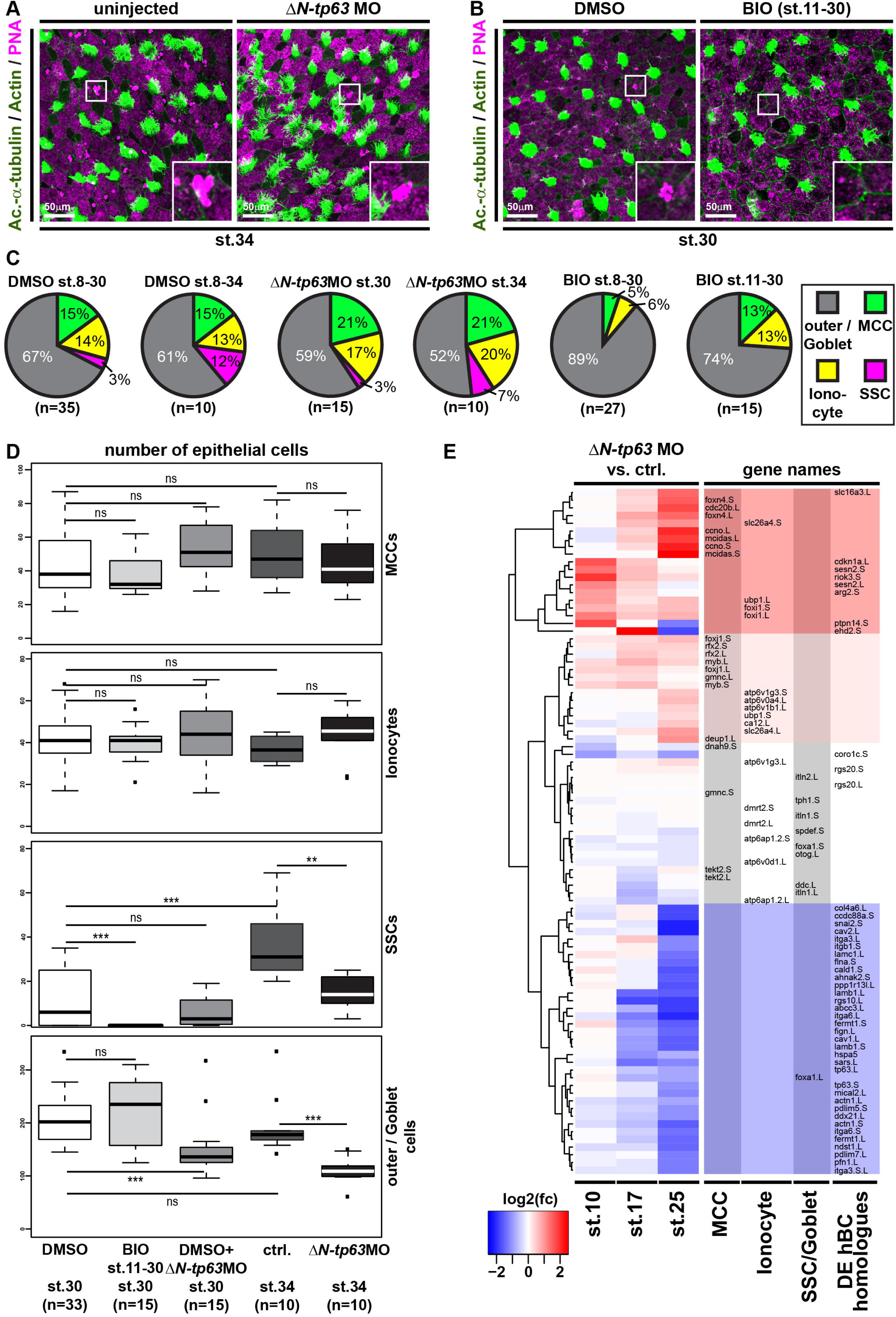
Knockdown of *ΔN-tp63* stimulates MCC and Ionocyte specification at the expense of BC and SSCs in *Xenopus*. **(A-D)** Analysis of cell type composition by confocal microscopy and staining for MCCs (Ac.-α-tubulin, green), Ionocytes (no staining, black), SSCs (large vesicles, PNA staining, magenta) and outer/Goblet cells (small granules, PNA staining, magenta) at st. 34 (A) and st. 30 (B). Actin staining (green). **(A)** *ΔN-tp63* MO increases MCC and Ionocyte numbers, but reduces numbers of SSCs. **(B)** BIO application from st. 11 does not affect MCC and Ionocyte numbers, but prevents specification of SSCs. **(C,D)** Quantification from (A,B). Mann Whitney test, not significant, ns = *P*>0.05; * = *P*≤ 0.01; *** = *P* 0.001. **(E)** RNA-sequencing at st.10, 17 and 25 on *Xenopus* mucociliary organoids comparing controls to *ΔN-tp63* MO injected. N=3 per stage and treatment. Heatmap and hierarchical clustering of log2 fold changes (fc) in cell type gene expression in *ΔN-tp63* morphants relative to controls. “Upregulated” clusters (red), “downregulated” cluster (blue). **(Related to Supplemental Figure S6)**

**Figure 5:**
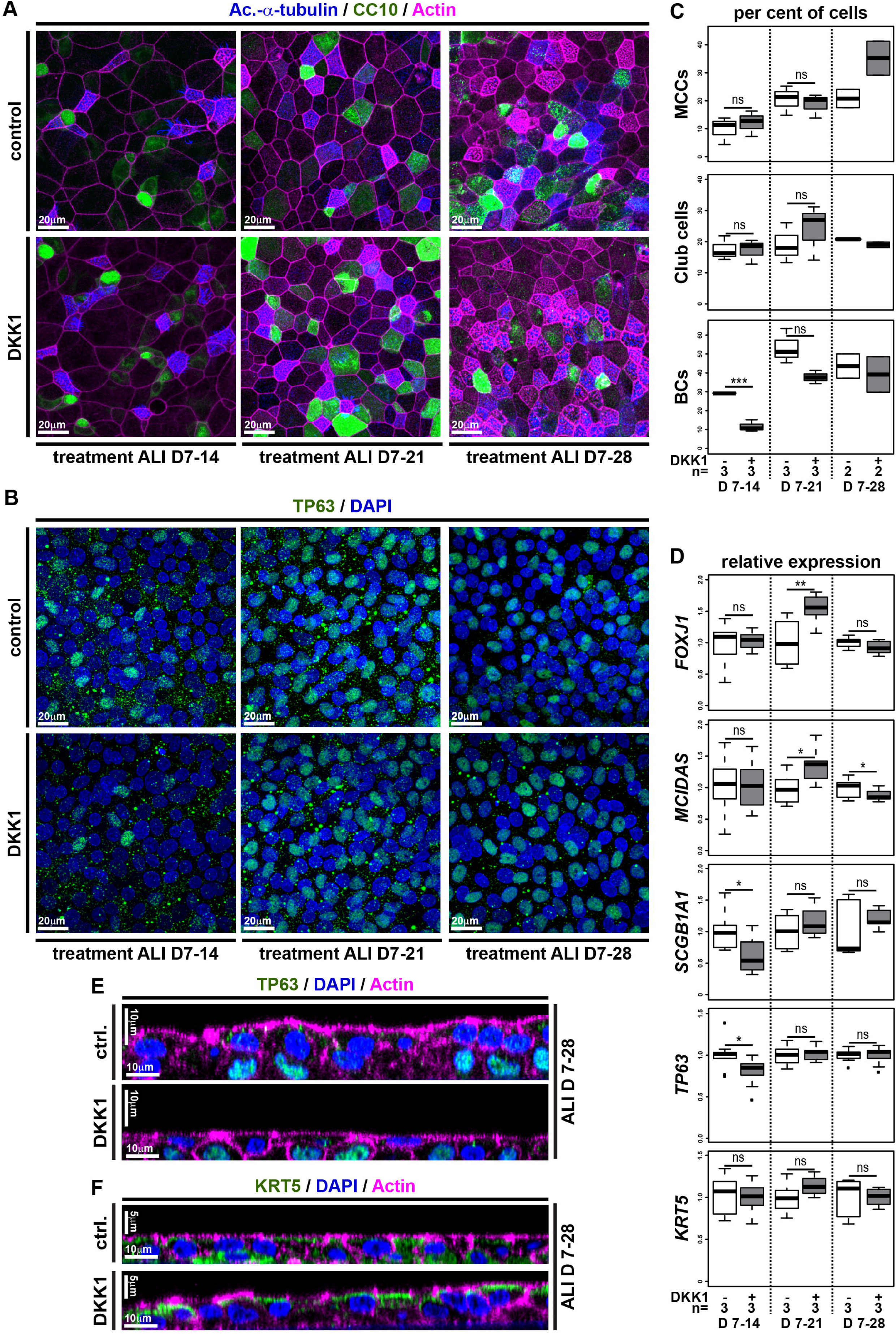
Inhibition of Wnt/β-catenin signaling transiently reduces stemness and promotes differentiation in human BCs. **(A-F)** BCIs in ALI culture for up to 4 weeks. Human recombinant DKK1 (DKK1) was used to inhibit Wnt/β-catenin signaling starting at ALI day 7 (D7). **(A)** Confocal imaging of specimens stained for Ac.-α-tubulin (MCCs, blue), CC10 (Club cells, green) and Actin (cell membranes, magenta) shows moderately increased MCC and Club cell numbers after DKK1 treatment. **(B)** DKK1 leads to a transient decrease in BCs, but does not lead to loss of TP63+ (green) cells. Nuclei (DAPI, blue). **(C)** Quantification from (A,B). Mann Whitney test, not significant, ns = *P*>0.05; *** = *P*≤0.001. **(D)** qPCR expression levels are depicted relative to stage controls. DKK1 increases expression of MCC (*FOXJ1*, *MCIDAS*) and to a lesser extent Club cell (*SCGB1A1*) markers, but without reduction of BC markers (*TP63*, *KRT5*). Student’s t-test, not significant, ns = *P*>0.05; * = *P*≤0.05; ** = *P*≤0.01. **(E,F)** Optical orthogonal sections of confocal images after staining for BC markers TP63 (green, in E) and KRT5 (green, in F). **(Related to Supplemental Figure S6)**

### Wnt/β-catenin-induced block of differentiation is reversible

Given the importance of correctly regulated Wnt/β-catenin signaling for BCs as well as for the generation of correct cell type-composition in mucociliary epithelia, we were interested to elucidate if prolonged exposure to elevated Wnt signaling would alter BC behavior rendering them incompetent to (re-)generate a normal mucociliary epithelium. To address that, we treated *Xenopus* embryos with BIO starting at st. 8, but removed the drug from the medium at various stages and assessed cell type composition by immunofluorescence and ISH. Treatment with BIO from st. 8 until st. 30 caused reduced MCC, Ionocyte and SSC numbers, which recovered after wash-out of the drug and subsequent regeneration until st. 33 **(Figure 6A and Supplementary Figure S7A)**. Similarly, treatment with BIO from st. 8 until st. 17 confirmed reduced expression of cell type specification markers as well as a thickening of the *ΔN-tp63* expressing sensorial layer. Removal of the drug at st. 17 and regeneration until st. 25 brought back the expression of epithelial cell type markers and reduced sensorial layer thickness and *ΔN-tp63* expression, close to normal levels **(Figure 6B and Supplementary Figure S7B-D)**. To test if this remarkable regenerative ability was limited to *Xenopus* development, we conducted analogous experiments in BCI ALI cultures. BCI cultures were treated with RSPO2 from ALI day 7 until day 21, resulting in deficient formation of MCCs and Club cells. Then, RSPO2 was removed from the medium and BCIs were allowed to recover for 7 days. After removal of RSPO2, BCIs were able to successfully regenerate MCCs and Club cells and to express cell type markers for epithelial cell types, without drastic changes in proliferation or epithelial cell numbers **(Figure 6C and Supplementary Figure S8A-E)**. Additionally, orthogonal optical sections of RSPO2-treated BCIs after recovery revealed a normalization of epithelial thickness and KRT5 staining **(Figure 6D,E)**. Collectively, our work revealed that excessive levels of Wnt signaling cause overactivation of ΔN-TP63 and block specification of epithelial cell types in a reversible manner, without altering the potential of BCs to form MCCs and secretory cells.

**Figure 6:**
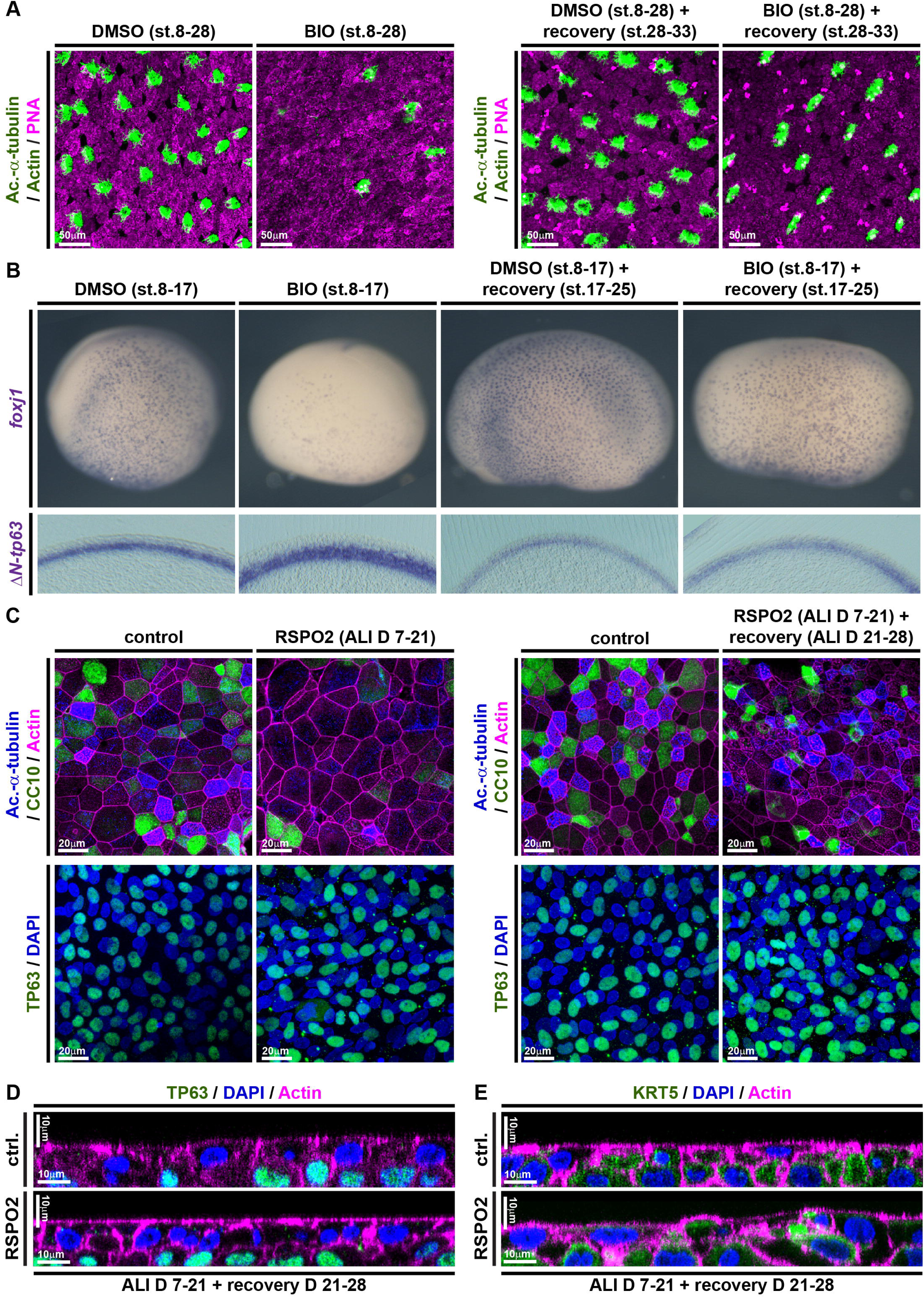
Wnt/β-catenin-induced increase in BCs and loss of epithelial differentiation are reversible. **(A,B)** In *Xenopus*, BIO treatments from st. 8-17 or st. 8-30 inhibit differentiation as compared to DMSO treated controls, but the epithelium can regenerate after removal of BIO and recovery until st. 33 (A) or st. 25 (B). **(A)** BIO treatment reduces MCC (Ac.-α-tubulin, green), Ionocyte (no staining, black), and SSC (large vesicles, PNA staining, magenta) numbers in confocal micrographs at st. 28, which recover after regeneration until st. 33. Actin staining (green). **(B)** ISH shows reduction in *foxj1* expressing cells and an increase in *ΔN-tp63* expression in BIO treated whole mounts (upper row) and transversal sections (bottom row) at st. 17, which both recover after regeneration until st. 25. DMSO (N=5), BIO st.17 (N=5), DMSO st.25 (N=5), BIO+recovery st.25 (N=5) **(C)** Confocal imaging of specimens stained for Ac.-α-tubulin (MCCs, blue), CC10 (Club cells, green) and Actin (cell membranes, magenta) show reduced MCC and Club cell numbers in RSPO2-treated cultures from ALI D7 – D21, but regeneration of MCCs and Club cells at ALI D28. N=3 cultures per timepoint and treatment (upper panels). No loss of TP63+ (green) BCs is observed in these experiments (bottom panels). Nuclei (DAPI, blue). **(D,E)** Optical orthogonal sections of confocal images. RSPO2-treated cultures display normalized epithelial thickness and staining for BC markers TP63 (green, in D) and KRT5 (green, in E) after regeneration at ALI D28. **(Related to Supplemental Figures S7 and S8)**

## Discussion

Our work demonstrates a requirement for dynamically regulated Wnt/β-catenin signaling in multiciliated (MCCs) and Basal (BCs) cells of the developing and regenerating mucociliary epithelium as well as a pro-proliferative effect of Wnt/β-catenin in mucociliary epithelia. Elevated Wnt/β-catenin signaling blocks cell fate specification of ciliated and secretory cells from BCs, while Wnt signaling during stages of differentiation promotes MCCs differentiation and ciliogenesis.

In BCs, high Wnt/β-catenin levels promote the expression of *ΔN-tp63*, a hallmark marker for BCs in various epithelia, including the mammalian respiratory tract (Hogan et al., 2014; Soares and Zhou, 2018; Zuo et al., 2015). We also provide evidence that *ΔN-tp63* is necessary and sufficient to promote BC fate and to inhibit specification into mature epithelial cells, including MCCs and secretory cells. *ΔN-tp63* was previously shown to be directly regulated by β-catenin in ChIP-seq studies in mammals and *Xenopus* and by promoter analysis (Kjolby and Harland, 2017; Ruptier et al., 2011). ΔN-tp63 is also known to inhibit differentiation and to promote proliferation in various cancers as well as in skin BCs (keratinocytes); in part this is regulated by transcription of cell cycle and pro-proliferative genes, which are also regulated by ΔN-tp63 in *Xenopus* (Chen et al., 2018; Soares and Zhou, 2018). Thus, our work provides a mechanistic explanation for the negative effects of elevated Wnt/β-catenin on mucociliary differentiation reported in multiple studies and implicates ΔN-tp63 as potential driver of proliferation after mucociliary injury. Interestingly, ΔN-tp63 is not required for initial formation of a mucociliary epithelium during development (Daniely et al., 2004). Nevertheless, we found that ΔN-tp63 expression is required for maintenance of BCs and that a loss of ΔN-tp63 leads to excessive cell fate specification and differentiation of MCCs and Ionocytes causing a deficiency in late-specified SSCs in the *Xenopus* epidermis, thereby, altering mucociliary cell type composition. Similar observations were made in developing *tp63^−/−^* mice, in which airway mucociliary epithelia formed during development, but those epithelia presented an excess of MCCs and a loss of BCs (Daniely et al., 2004). Furthermore, knockdown of *TP63* in ALI cultures of human airway cells prevents regeneration and causes senescence, indicating loss of stemness (Arason et al., 2014). Collectively, these findings support the conclusion that ΔN-tp63 is a Wnt/β-catenin-regulated master transcription factor in BCs, deciding between BC maintenance and differentiation. Interestingly, Wnt/β-catenin is dispensable to maintain ΔN-TP63 and BCs after epithelialization in regenerating BCIs, suggesting that *ΔN-TP63* can be maintained by other pathways when BCs are confluent. This could be achieved by Notch signaling, which was previously shown to regulate BCs (Rock et al., 2011), but requires cell-cell contact provided by a sufficient density of cells. Furthermore, our data demonstrate a deep evolutionary conservation of signaling and regulatory mechanisms across vertebrate mucociliary epithelia, establish the *Xenopus* epidermis as a new model to study BCs *in vivo*, and provide a set of conserved BC genes, which can be used as BC markers in *Xenopus* and studied mechanistically in the future.

In MCCs, Wnt/β-catenin signaling during differentiation stages is required for normal ciliation. These results are in line with previous work demonstrating that Wnt/β-catenin signaling is necessary for normal ciliogenesis in various vertebrate systems by co-regulating *foxj1*, a master transcription factor for motile cilia (Caron et al., 2012; Sun et al., 2019; Walentek et al., 2015). The positive effect of Wnt/β-catenin on MCC specification suggested by some studies could be explained by the extensive positive cross-regulation between transcription factors expressed in MCCs. The multiciliogenesis cascade is initiated by a transcriptional regulatory complex consisting of Multicilin (encoded by *mcidas*), E2f4/5 and TfDp1, which activates expression of the downstream transcription factors *foxj1*, *rfx2/3*, *myb*, *tp73* and *foxn4* (Quigley and Kintner, 2017; Stubbs et al., 2012; Walentek and Quigley, 2017). These transcription factors generate a positive feedback on their expression (Choksi et al., 2014; Quigley and Kintner, 2017). This positive cross-regulation was especially well demonstrated for FOXJ1 and RFX2/3, and argues for the possibility that activation of *FOXJ1* could ultimately lead to activation of the multiciliogenesis cascade (Didon et al., 2013). Additionally, we have previously found that *myb* expression is downregulated after inhibition of Wnt/β-catenin signaling, suggesting that *myb* could be regulated by Wnt signaling as well (Tan et al., 2013; Walentek et al., 2015). Thus, different levels and timing of Wnt/β-catenin signaling activation could provide an explanation as to why some studies reported negative effects on MCC formation, while others described an increase in MCC numbers.

Our data argue that the loss of differentiated MCCs upon excessive Wnt activation is a consequence of impaired specification, rather than Goblet cell hyperplasia as previously suggested (Mucenski et al., 2005). While we did not observe an increase in MUC5B expressing cells in the epithelium after Wnt activation, we did detect elevated *MUC5B* expression levels. This suggests potential induction of subepithelial Goblet cell formation *in vitro* after overactivation of Wnt/β-catenin signaling, similar to the induction of Goblet cells in submucosal glands (Driskell, 2004).

Finally, our data indicate that persistent Wnt/β-catenin activation in mucociliary epithelia could lead to BC hyperplasia and a remodeling of the epithelium. Importantly, we show that these effects are reversible and that a return to normal signaling levels can promote re-establishment of a differentiated epithelium. This is an important notion in the context of chronic lung diseases, such as COPD, which are also associated with defective epithelial differentiation, BC hyperplasia, altered Wnt ligand expression, and overactivation of the Wnt/β-catenin pathway (Baarsma and Königshoff, 2017; Chen et al., 2010; Heijink et al., 2013; Königshoff et al., 2008). Furthermore, it was shown that nasal polyps from chronic rhinosinusitis patients produce excess levels of WNT3a and MCC differentiation is inhibited, but that MCC formation could be rescued by application of a Wnt-inhibitor (Dobzanski et al., 2018). Together, these findings suggest that even in a chronically pathogenic state, targeted Wnt/β-catenin signaling inhibition could provide a potential avenue for treatment of patients with COPD and other chronic lung diseases, for which treatment options are currently limited or absent.

## Acknowledgements

We thank: S. Schefold, D. and E. Pangilinan, J. Groth for technical help; R. Kjolby, W. Finkbeiner, C. Boecking, G. Pyrowolakis, L. Kodjabachian, R. Harland, B. Gomperts for discussions and sharing of unpublished results; R. Crystal and M. Walters for BCi-NS1.1 cells; T. Kwon and S. Medina Ruiz for genome annotation files; NXR (RRID:SCR_013731), Xenbase (RRID:SCR_004337), EXRC for Xenopus resources; Transcriptome and Genome Analysis laboratory Göttingen for deep sequencing; Light Imaging Center Freiburg for microscope use; JAX (RRID:SCR_004633) for mice; Galaxy Europe for bioinformatics platform.

This study was supported by the German Research Foundation (DFG) under the Emmy Noether Programme (grant WA3365/2-1) and under Germany’s Excellence Strategy (CIBSS – EXC-2189 – Project ID 390939984) as well as by an NHLBI Pathway to Independence Award (K99HL127275) to PW. Generation of transgenic Xenopus was supported by FWO-Vlaanderen grant G029413N and by the Concerted Research Actions from Ghent University (BOF15/GOA/011) to KV. DIS was supported by the URAP program and a Pergo Foundation SURF L&S Fellowship.

## Author contributions

MH, DIS, MB, AT, PW: Xenopus experiments.

JLGV, AT, PW: tissue culture experiments.

JLGV, PW: mouse Wnt-reporter analysis.

HTT, KV: generated Xenopus laevis Wnt reporter line.

OS, PW: bioinformatics.

MH, PW: experimental design and planning.

PW: study design and supervision, coordinating collaborative work, manuscript preparation.

## STAR Methods

### Contact for reagent and resource sharing

Further information and requests for resources and reagents should be directed to and will be fulfilled by the Lead Contact, Peter Walentek.

Immortalized human Basal cells (BCIs) were generated and are distributed by the Crystal laboratory at Genetic Medicine/Joan and Sanford I. Weill Department of Medicine, Weill Cornell Medical School, New York, USA. Sharing of this resource is subject to an MTA.

The Wnt reporter line *Xla.Tg(WntREs:dEGFP)^Vlemx^* was obtained from the National Xenopus Resource (NXR) at Marine Biological Laboratory, Woods Hole, USA, and the European Xenopus Resource Centre (EXRC) at University of Portsmouth, School of Biological Sciences, UK. Sharing of this resource is subject to an MTA.

### Experimental models and subject details

#### Xenopus laevis

Wildtype and transgenic *Xenopus laevis* were obtained from the National Xenopus Resource (NXR) at Marine Biological Laboratory, Woods Hole, USA, and the European Xenopus Resource Centre (EXRC) at University of Portsmouth, School of Biological Sciences, UK. Frog maintenance and care was conducted according to standard procedures and based on recommendations provided by the international Xenopus community resource centers NXR and EXRC as well as by Xenbase (xenbase.org). All experiments were conducted in embryos derived from at least two different females and independent in vitro fertilizations.

#### Mice

Mice from the strain TCF/Lef1-HISTH2BB/EGFP (61Hadj/J) (Ferrer-Vaquer et al., 2010) were obtained from the Jackson Laboratories (JAX) and genotyped using the protocol deposited under jax.org/strain/013752. Reporter analysis was conducted on tissues derived from male and female animals and no differences were observed between the sexes. Animal care was conduced by centralized facilities and according to standard procedures.

#### Immortalized human Basal cells (BCIs)

BCIs were generated as described in (Walters et al., 2013) and were provided by the Crystal laboratory. All experiments were conducted on cells derived from the same passage (passage 10). Expansion and ALI cultures of BCIs were conducted according to (Walters et al., 2013) at 37 °C.

#### Ethics statements on animal experiments

This work was done in compliance with German animal protection laws and was approved under Registrier-Nr. X-18/02F and G-18/76 by the state of Baden-Württemberg, as well as with approval of University of California, Berkeley’s Animal Care and Use Committee. University of California Berkeley’s assurance number is A3084-01, and is on file at the National Institutes of Health Office of Laboratory Animal Welfare.

### Method details

#### Manipulation of *Xenopus* Embryos, Constructs and In Situ Hybridization

*X. laevis* eggs were collected and *in vitro*-fertilized, then cultured and microinjected by standard procedures (Sive et al., 2010). Embryos were injected with Morpholino oligonucleotides (MOs, Gene Tools) and mRNAs at the four-cell stage using a PicoSpritzer setup in 1/3x Modified Frog Ringer’s solution (MR) with 2.5% Ficoll PM 400 (GE Healthcare, #17-0300-50), and were transferred after injection into 1/3x MR containing Gentamycin. Drop size was calibrated to about 7–8 nL per injection.

Morpholino oligonucleotides (MOs) were obtained from Gene Tools targeting *ctnnb1.L and .S* (Heasman et al., 2000), or targeting *ΔN-tp63.L and .S* (this study), and used at doses ranging between 34 and 51ng (or 4-6pmol).

*ΔN-tp63* was cloned from total cDNA into pCS107 using ΔN-tp63-f and ΔN-tp63-stop-r primers matching NCBI reference sequence XM_018261616.1. BamH1/Sal1 restriction enzymes were used for subcloning. A *gfp-ΔN-tp63* fusion construct was generated using gfp-f and gfp-r, and BamH1 restriction enzyme to fuse GFP to the N-terminus of pCS107-ΔN-tp63. A hormone inducible *gfp-ΔN-tp63-gr* fusion construct was generated using anon stop-sequence (ΔN-tp63-nonstop-r), primers for the GR-domain (Kolm and Sive, 1995) gr-lbd-f and gr-lbd-r, and Sal1 restriction enzymes to fuse the GR domain to the C-terminus of pCS107-gfp-ΔN-tp63. All sequences were verified by Sanger sequencing, and linearized with Asc1 to generate mRNAs (used at 150ng/μl) and pCS107-*ΔN-tp63* with BamH1 to generate an anti-sense probe template.

mRNAs encoding membrane-GFP or membrane-RFP or Centrin4-CFP (Antoniades et al., 2014) were used in some experiments as lineage tracers at 50 ng/µL (not shown). All mRNAs were prepared using the Ambion mMessage Machine kit using Sp6 (#AM1340).

DNAs were purified using the PureYield Midiprep kit (Promega, #A2495) and were linearized before in vitro synthesis of anti-sense RNA probes using T7 or Sp6 polymerase (Promega, #P2077 and #P108G), RNAse Inhibitor (Promega #N251B) and Dig-labeled rNTPs (Roche, #3359247910 and 11277057001). Embryos were *in situ* hybridized according to (Harland, 1991), bleached after staining and imaged. Sections were made after embedding in gelatin-albumin with glutaraldehyde at 50-70μm as described in (Walentek et al., 2012).

Drug treatment of embryos started and ended at the indicated stages. DMSO (Sigma, #D2650) or ultrapure Ethanol (NeoFroxx #LC-8657.3) were added to the medium as vehicle controls. 6-Bromoindirubin-3’-oxime (BIO, Sigma-Aldrich/Merck #B1686) was used in DMSO at 75 μM (BIO low) or 150 μM (BIO high). Dexamethasone (Sigma-Aldrich/Merck #D4902) was used in Ethanol at 10μM.

Morpholino nucleotide and cloning primer sequences:

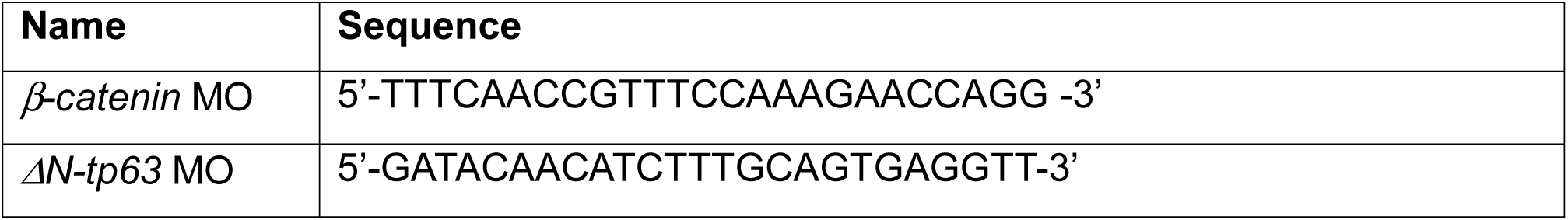
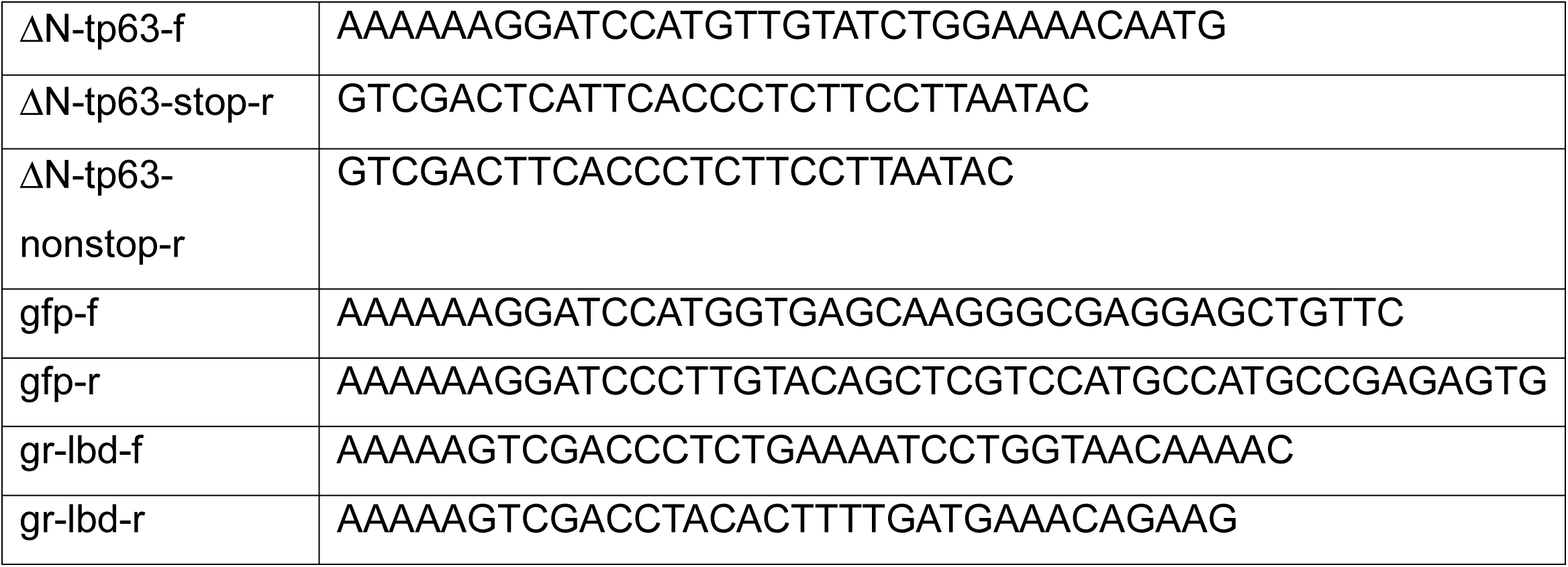

#### Generation of the *Xenopus* Wnt/β-catenin signaling reporter line

The Wnt reporter line *Xla.Tg(WntREs:dEGFP)^Vlemx^* was generated using the sperm nuclear transfer method as described in detail in (Hirsch et al., 2002). The Wnt-responsive promoter consists of 7 copies of a TCF/LEF1 binding DNA element and a minimal TATA box and a reporter gene encoding destabilized EGFP and a polyA sequence. The transgene is flanked on both sides by two copies of the chicken HS4-core sequence (Tran et al., 2010).

#### RNA-sequencing on *Xenopus* mucociliary organoids and bioinformatics analysis

*X. laevis* embryos were either injected 4x into the animal hemisphere at four-cell stage with *ΔN-tp63* MO or remained uninjected, and were cultured until st. 8. Animal caps were dissected in 1x Modified Barth’s solution (MBS) and transferred to 0.5x MBS + Gentamycin (Sive et al., 2010). 10-15 organoids were collected in TRIzol (Thermo Fisher #15596026) per stage at st. 10.5 (st. 10), st. 16-19 (st. 17) and st. 24-25 (st. 25). Organoids were derived from 3 independent experiments.

500 ng total RNA per sample was used, poly-A selection and RNA-sequencing library preparation was done using non strand massively-parallel cDNA sequencing (mRNA-Seq) protocol from Illumina, the TruSeq RNA Library Preparation Kit v2, Set A (Illumina #RS-122-2301) according to manufacturer’s recommendation. Quality and integrity of RNA was assessed with the Fragment Analyzer from Advanced Analytical by using the standard sensitivity RNA Analysis Kit (Advanced Analytical #DNF-471). All samples selected for sequencing exhibited an RNA integrity number over 8. For accurate quantitation of cDNA libraries, the QuantiFluor™dsDNA System from Promega was used. The size of final cDNA libraries was determined using the dsDNA 905 Reagent Kit (Advanced Bioanalytical #DNF-905) exhibiting a sizing of 300 bp on average. Libraries were pooled and paired-end 100bp sequencing on a HiSeq2500 was conducted at the Transcriptome and Genome Analysis Laboratory, University of Göttingen. Sequence images were transformed with Illumina software BaseCaller to BCL files, which was demultiplexed to fastq files with bcl2fastq v2.17.1.14. Quality control was done using FastQC v0.11.5 (Andrews, Simon (2010). “FastQC a quality-control tool for high-throughput sequence data” available at http://www.bioinformatics.babraham.ac.uk/projects/fastqc).

Sequencing generated a total of 2x 581.7 Mio reads (average 2x 32.3 Mio / library). After adapter-trimming, paired-end reads were mapped to *Xenopus laevis* genome assembly v9.2 using RNA STAR v2.6.0b-1 (Dobin et al., 2013). featureCounts v1.6.3 (Liao et al., 2014) was used to count uniquely mapped reads per gene and statistical analysis of differential gene expression was conducted in DEseq2 v1.22.1 (Love et al., 2014). Go-term analysis was done with “humanized” versions of *Xenopus* gene names (by removing “.L” and “.S” from the name) using the GO Consortium website (geneontology.org). Heatmaps were generated in R v3.5.1 using ggplot2/heatmap2 v2.2.1. All bioinformatic analysis was performed on the Galaxy / Europe platform (usegalaxy.eu).

#### Air-liquid interface (ALI) culture of immortalized human Basal cells (BCIs)

ALI cultures of BCIs were conducted according to (Walters et al., 2013) on Costar Transwell Filters (Costar #3470), coated with human Type IV Collagen (Sigma #C7521) dissolved in Acetic acid (Carl Roth #3738.4). For Basal cell expansion the BEGM Bullet Kit (Lonza #CC-3170) was used with all supplements as recommended by the manufacturer, but without the antibiotics. Instead Penicillin-Streptomycin (0.5%, Sigma #P4333), Gentamycin sulphate (0.1%, Carl Roth, #2475.1) and Amphotericin B (0.5%, Gibco #15290-018) were added. Differentiation of cells was conducted in DMEM:F12 (Gibco #11330-032) with UltroserG (2%, Pall BioSphera-Science #15950-017 dissolved in sterile cell culture grade water from Gibco #15230-071), and Penicillin-Streptomycin (0.5%, Sigma #P4333), Gentamycin sulphate (0.1%, Carl Roth, #2475.1) and Amphotericin B (0.5%, Gibco #15290-018) for up to 28 days. Media were filtered (0.22 μm) before use. Manipulations of Wnt signaling were done by addition of human recombinant RSPO2 (R&D systems 3266-RS) or human recombinant DKK1 (R&D systems 5439-DK), which were reconstituted in sterile PBS, pH 7.4 containing 0.1% bovine serum albumin at 200ng/ml.

#### ALI culture of primary mouse tracheal epithelial cells (MTECs)

ALI cultures of MTECs were conducted according to (Vladar and Brody, 2013) on Costar Transwell Filters (Costar #3470), coated with rat tail Collagen (BD Biosciences #354236) in Acetic acid. Cells were isolated from TCF/Lef1-HISTH2BB/EGFP (61Hadj/J). The following reagents and supplements were used as indicated in the protocol:

| Name | Vendor & P/N | Final concentration | Medium |
| --- | --- | --- | --- |
| Pronase | Roche<br>#10165921001 | 1.5mg/ml | Pronase solution |
| Ham's F12 | Life Technologies<br>#11765054 | - | Pronase solution,<br>DNAse solution |
| DNAse | Sigma #DN25 | 0.5mg/ml | DNAse solution |
| DMEM:F12 | Gibco #11330-032 | - | Proliferation Medium,<br>Differentiation |
|  |  |  | Medium |
| Penicillin-Streptomycin | Sigma #P4333 | 1% | Proliferation Medium,<br>Differentiation<br>Medium,<br>Pronase solution |
| Amphotericin B | Gibco #15290-018 | 0.1% | Proliferation Medium,<br>Differentiation<br>Medium |
| Sodium Biscarbonate | Life Technologies<br>#25080060 | 0.3% | Proliferation Medium,<br>Differentiation<br>Medium |
| Insulin | Sigma #1182 | 10µg/ml | Proliferation Medium |
| Epidermal Growth Factor | BD Biosciences<br>#354001 | 25µg/ml | Proliferation Medium |
| Apo-Transferrin | Sigma #T1147 | 5µg/ml | Proliferation Medium |
| Cholera toxin | Sigma #C8052 | 0.1µg/ml | Proliferation Medium |
| Bovine pituitary extract | Sigma #SLBV9702 | 30µg/ml | Proliferation Medium |
| FBS Superior | Biochrom #S0615 | 5% | Proliferation Medium |
| Retinoic acid | Sigma #R2625 | 50nM | Proliferation Medium,<br>Differentiation<br>Medium |
| NuSerum | BD Biosciences<br>#355100 | 2% | Differentiation<br>Medium |

Primaria cell culture dishes (Corning #353803) were used for selection during the procedure. Cells were cultured for up to 21 days.

#### Quantitative RT-PCR on cDNAs from BCIs

Before total RNA extraction from BCIs, filters were washed 3x 5 min with PBS and removed from the insets using a scalpel cleaned with RNase away (MbP #7002). The RNeasy Mini Kit (Qiagen #74104) was used, and the cells were collected in RLT buffer + β-Mercaptoethanol (10 μl / ml), votexted for 2 min, and lysed using QIAshredder (Qiagen #79654) columns. RNA was collected in UltraPure water (Invitrogen #10977-035) and used for cDNA synthesis with iScript cDNA Synthesis Kit (Bio-Rad #1708891). qPCR-reactions were conducted using Sso Advanced Universal SYBR Green Supermix (Bio-Rad #172-5275) on a CFX Connect Real-Time System (Bio-Rad) in 96-well PCR plates (Brand #781366). Experiments were conducted in biological and technical triplicates and normalized by *GAPDH* and *ODC* expression levels. Expression levels were analyzed in Excell and graphs were generated using R.

Primers:

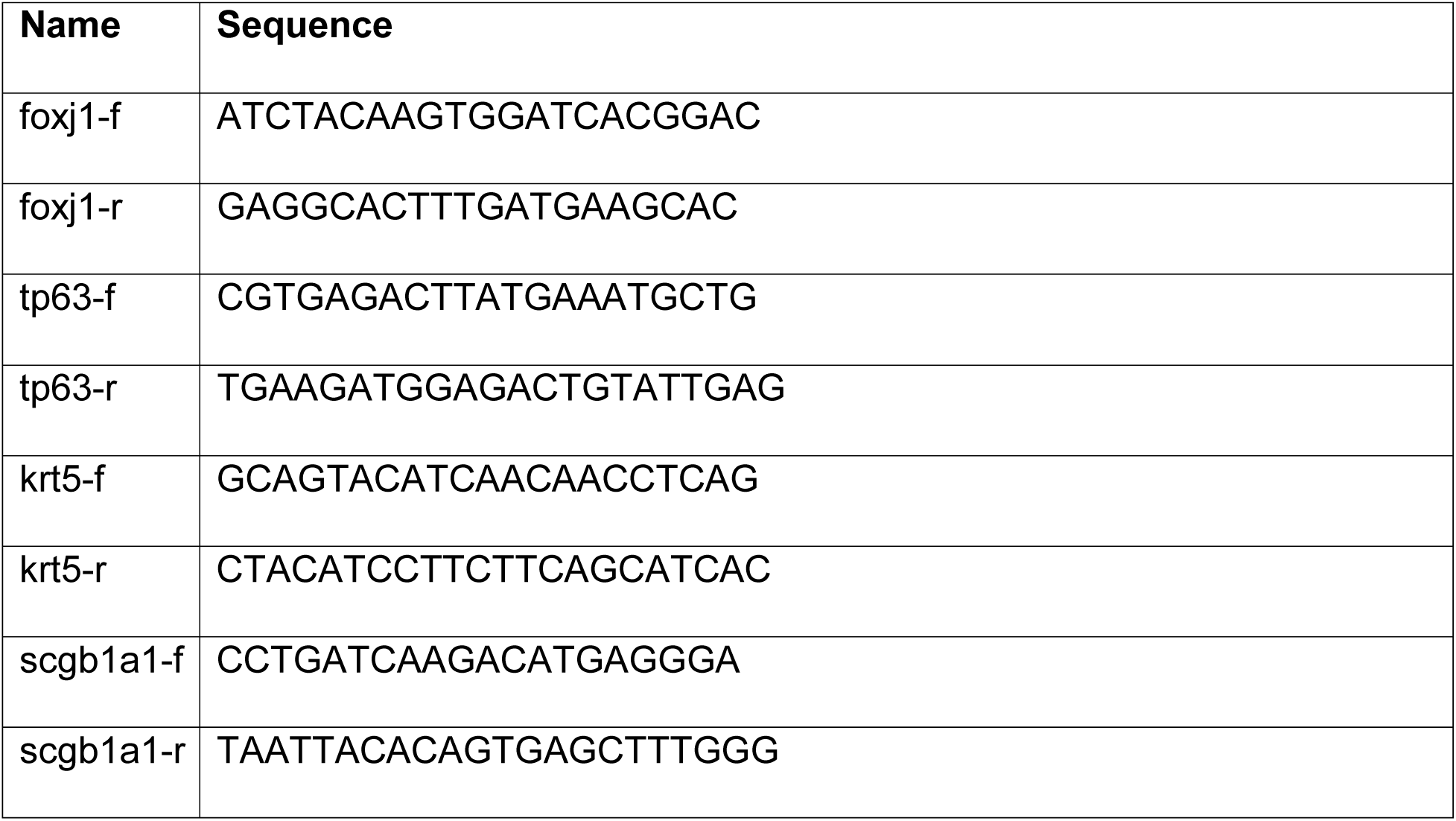
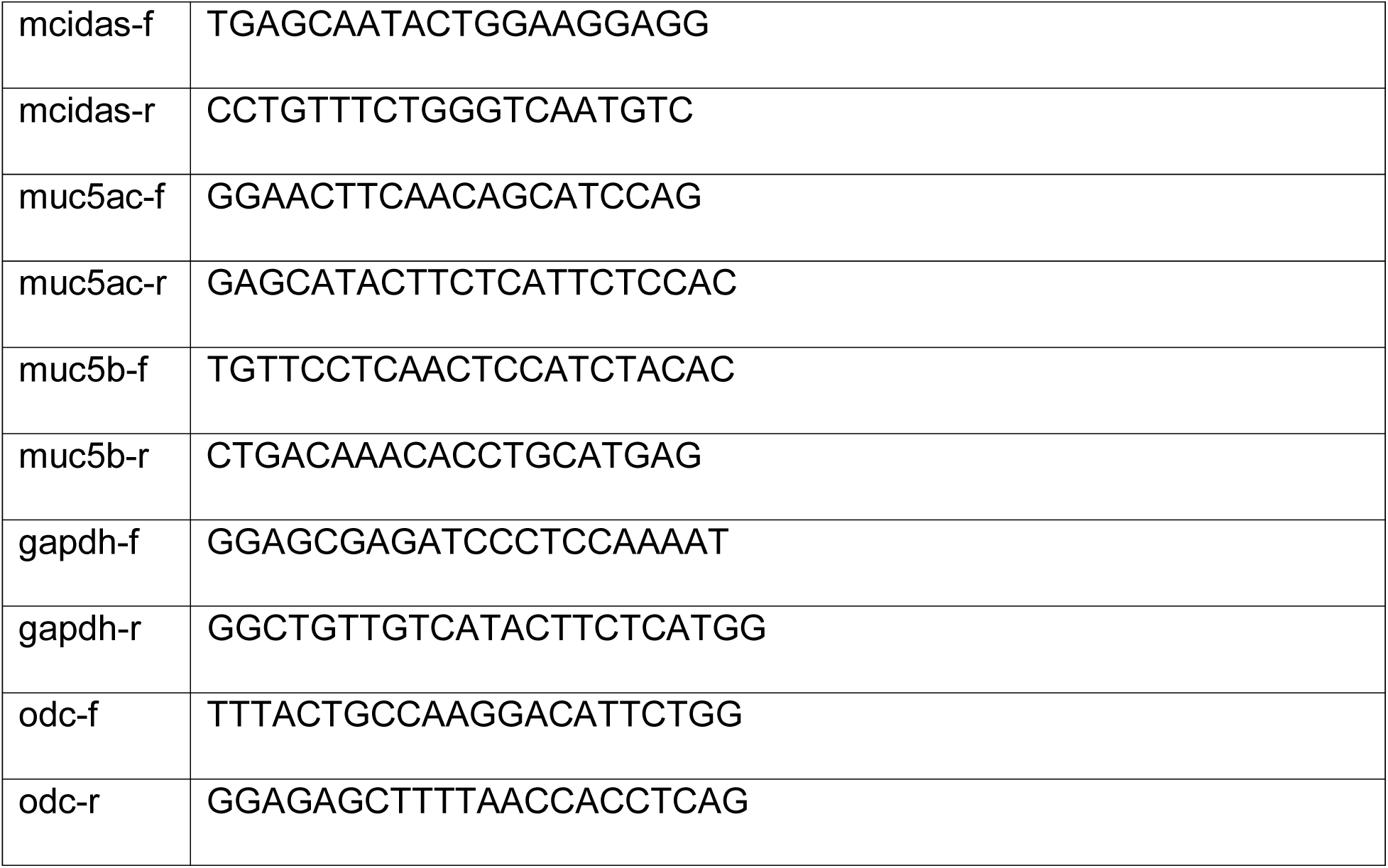

#### Immunofluorescence Staining and Sample Preparation

Whole *Xenopus* embryos, were fixed at indicated stages in 4% paraformaldehyde at 4 °C over-night or 2 h at room temperature, then washed 3x 15 min with PBS, 2x 30 min in PBST (0.1% Triton X-100 in PBS), and were blocked in PBST-CAS (90% PBS containing 0.1% Triton X-100, 10% CAS Blocking; ThermoFischer #00-8120) for 1h at RT. For cryo sections, embryos were equilibrated in 50% Sucrose at 4 °C over-night, embedded in O.C.T. cryomedium (Tissue-Tek #25608-930), frozen at −80 °C, and sectioned at 30-50μm. Immunostaining on sections was done as for whole embryos after initial 3x 15 min washes with PBS and 15 min re-fixation in 4% paraformaldehyde at RT.

Mouse lungs were dissected, washed in ice-cold PBS several times and fixed at indicated stages in 4% paraformaldehyde at 4 °C for >24 h. The tissue was then equilibrated in 50% Sucrose at 4 °C over-night, embedded in O.C.T. cryomedium (Tissue-Tek #25608-930), frozen at −80 °C, and sectioned at 10-14 μm. For immunostaining, sections were washed 3x 15 min with PBS and re-fixed in 4% paraformaldehyde at RT 15 min followed by 2x 30 min washes in PBST (0.1% Triton X-100 in PBS). Samples were blocked in PBST-CAS (90% PBS containing 0.1% Triton X-100, 10% CAS Blocking) for 30 min – 1 h at RT.

MTEC and BCI cells grown in ALI culture were washed 3x 5 min with PBS before fixation in 4% paraformaldehyde at 4 °C for >24 h. The culture filters were removed from the insets using a scalpel, divided into multiple parts and used for different combinations of stains. Filter parts were washed 2x 30 min in PBST (0.1% Triton X-100 in PBS) and blocked in PBST-CAS (90% PBS containing 0.1% Triton X-100, 10% CAS Blocking) for 30 min – 1 h at RT.

Primary antibodies used in this study:

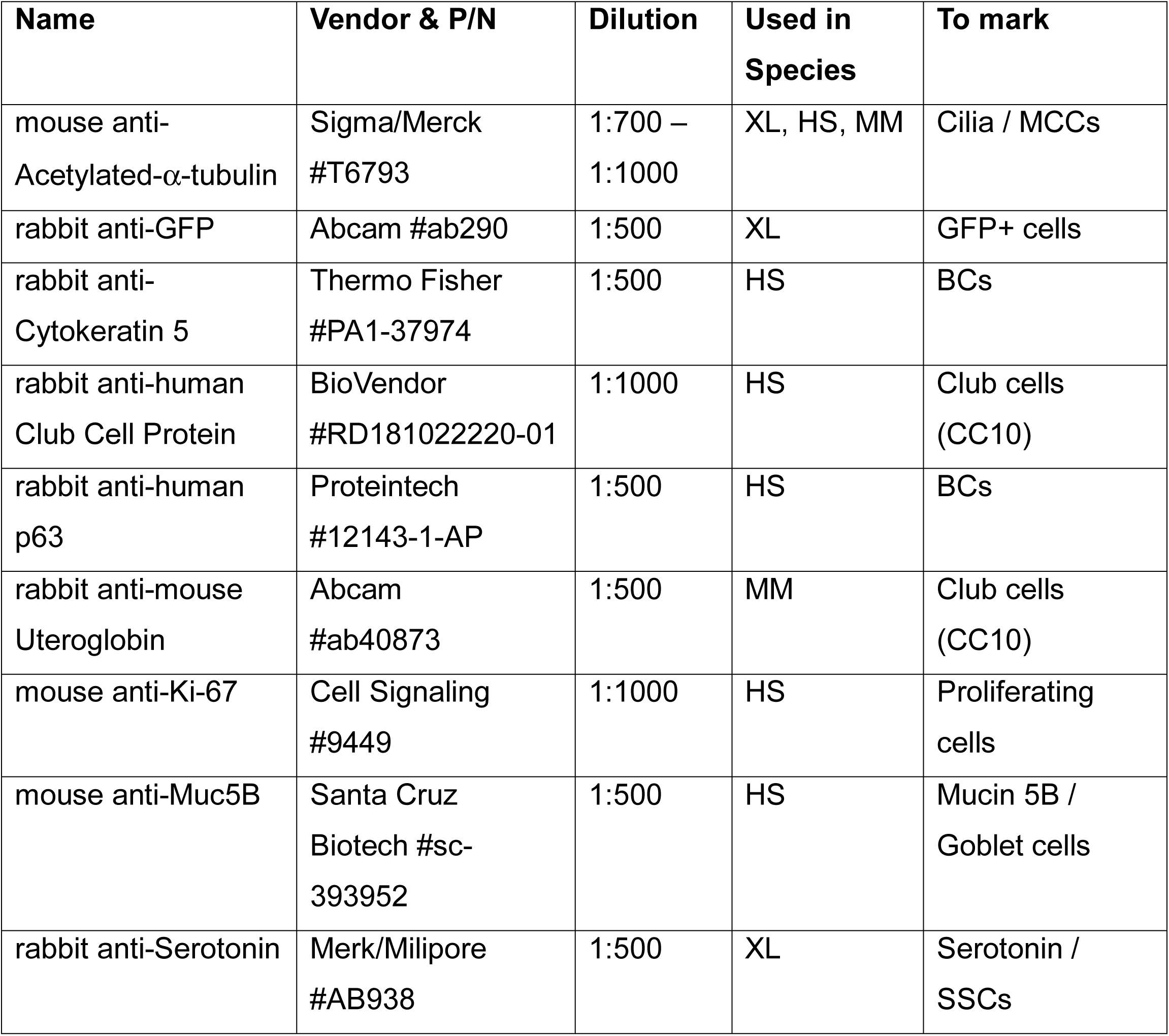

| Name | Vendor & P/N | Dilution | Used in Species | To mark |
| --- | --- | --- | --- | --- |
| mouse anti-Acetylated-α-tubulin | Sigma/Merck #T6793 | 1:700 – 1:1000 | XL, HS, MM | Cilia / MCCs |
| rabbit anti-GFP | Abcam #ab290 | 1:500 | XL | GFP+ cells |
| rabbit anti-Cytokeratin 5 | Thermo Fisher #PA1-37974 | 1:500 | HS | BCs |
| rabbit anti-human Club Cell Protein | BioVendor #RD181022220-01 | 1:1000 | HS | Club cells (CC10) |
| rabbit anti-human p63 | Proteintech #12143-1-AP | 1:500 | HS | BCs |
| rabbit anti-mouse Uteroglobin | Abcam #ab40873 | 1:500 | MM | Club cells (CC10) |
| mouse anti-Ki-67 | Cell Signaling #9449 | 1:1000 | HS | Proliferating cells |
| mouse anti-Muc5B | Santa Cruz Biotech #sc-393952 | 1:500 | HS | Mucin 5B / Goblet cells |
| rabbit anti-Serotonin | Merk/Milipore #AB938 | 1:500 | XL | Serotonin / SSCs |

Secondary antibodies (used at 1:250): AlexaFluor 555-labeled goat anti-mouse antibody (Molecular Probes #A21422), AlexaFlour 488-labeled goat anti-rabbit antibody (Molecular Probes #R37116), AlexaFlour 488-labeled donkey anti-mouse antibody (Molecular Probes #R37114), AlexaFlour 405-labeled goat anti-mouse antibody (Molecular Probes #A-31553). All antibodies were applied in 100% CAS Blocking (ThermoFischer #00-8120) over night at 4 °C or 2 h at RT (for secondary antibodies). DAPI was used to label nuclei (applied for 30 min. at room temperature, 1:100 in PBSt; Molecular Probes #D1306) in *Xenopus*. ProLong Gold Antifade Mountant with DAPI (Molecular Probes #P36931) was used to label nuclei in mouse and human samples. Actin was stained by incubation (30-120 min at room temperature) with AlexaFluor 488- or 647-labeled Phalloidin (1:40 in PBSt; Molecular Probes #A12379 and #A22287), mucus-like compounds in *Xenopus* were stained by incubation (over night at 4°C) with AlexaFluor 647-labeled PNA (1:1000 in PBSt; Molecular Probes #L32460).

#### Confocal imaging, image processing and analysis

Confocal imaging was conducted using a Zeiss LSM700 or Zeiss LSM880 and Zeiss Zen software. Wnt reporter sections from *Xenopus* and mice were imaged using tile-scans and images were reconstructed in ImageJ or Adobe Photoshop. Confocal images were adjusted for channel brightness/contrast and Z-stack projections or orthogonal sections were generated using ImageJ. A detailed protocol for quantification of *Xenopus* epidermal cell types was published (Walentek, 2018). Images of embryos after *in situ* hybridization and corresponding sections were imaged using an AxioZoom setup or AxioImager.Z1, and images were adjusted for color balance, brightness and contrast using Adobe Photoshop. Measurement of *ΔN-tp63* domain thickness in *Xenopus* was done in ImageJ using the NeuronJ plugin.

### Quantification and statistical analysis

#### Statistical Evaluation

Stacked bar graphs were generated in Microsoft Excel, box plots and heatmaps were generated in R (the line represents the median; 50% of values are represented by the box; 95% of values are represented within whiskers; values beyond 95% are depicted as outliers). Statistical evaluation of experimental data was performed using chi-squared tests (http://www.physics.csbsju.edu/stats/contingency.html), Wilcoxon sum of ranks (Mann-Whitney) tests (http://astatsa.com/WilcoxonTest/), or Student’s t-test (http://www.physics.csbsju.edu/stats/t-test.html) as indicated in figure legends.

#### Sample size and analysis

Sample sizes for all experiments were chosen based on previous experience and used embryos derived from at least two different females in *Xenopus*. Analysis of mouse Wnt-reporter was conducted in samples from N > 3 animals. No randomization or blinding was applied.

#### Use of shared controls

For parts of cell type quantification in *Xenopus* and BCIs, and qPCR experiments in BCIs shared controls or other conditions were used in multiple figures/graphs. Therefore, a detailed log of manipulation experiments in *Xenopus* and BCIs is provided in Supplemental Tabe 2. It contains information on experiment number, species/model, type of experiment, conditions, number of specimens, and in which figure/graph the data was used throughout the manuscript.

### Data and software availability

RNA-seq data have been deposited in the NCBI Gene expression Omnibus (GEO) database under the ID code: *AWAITING GEO ACCESSION NUMBER*.

## References

Antoniades, I., Stylianou, P., and Skourides, P.A. (2014). Making the Connection: Ciliary Adhesion Complexes Anchor Basal Bodies to the Actin Cytoskeleton. Dev. Cell 28, 70–80.

Arason, A.J., Jonsdottir, H.R., Halldorsson, S., Benediktsdottir, B.E., Bergthorsson, J.T., Ingthorsson, S., Baldursson, O., Sinha, S., Gudjonsson, T., and Magnusson, M.K. (2014). DeltaNp63 has a role in maintaining epithelial integrity in airway epithelium. PLoS One 9.

Baarsma, H.A., and Königshoff, M. (2017). ‘WNT-er is coming’L chronic lung diseases. Thorax 72, 746–759.

Borday, C., Parain, K., Thi Tran, H., Vleminckx, K., Perron, M., and Monsoro-Burq, A.H. (2018). An atlas of Wnt activity during embryogenesis in Xenopus tropicalis. PLoS One 13, e0193606.

Caron, A., Xu, X., and Lin, X. (2012). Wnt/ -catenin signaling directly regulates Foxj1 expression and ciliogenesis in zebrafish Kupffer’s vesicle. Development 139, 514–524.

Chen, Y., Peng, Y., Fan, S., Li, Y., Xiao, Z.X., and Li, C. (2018). A double dealing tale of p63: an oncogene or a tumor suppressor. Cell. Mol. Life Sci. 75, 965–973.

Chen, Y.T., Gallup, M., Nikulina, K., Lazarev, S., Zlock, L., Finkbeiner, W., and McNamara, N. (2010). Cigarette smoke induces epidermal growth factor receptor-dependent redistribution of apical MUC1 and junctional β airway epithelial cells. Am. J. Pathol. 177, 1255–1264.

Choksi, S.P., Lauter, G., Swoboda, P., and Roy, S. (2014). Switching on cilia: transcriptional networks regulating ciliogenesis. Development 141, 1427–1441.

Cibois, M., Luxardi, G., Chevalier, B., Thome, V., Mercey, O., Zaragosi, L.-E., Barbry, P., Pasini, A., Marcet, B., and Kodjabachian, L. (2015). BMP signalling controls the construction of vertebrate mucociliary epithelia. Development 142, 2352–2363.

Clevers, H. (2006). Wnt/β-Catenin Signaling in Development and Disease. Cell 127, 469–480.

Daniely, Y., Liao, G., Dixon, D., Linnoila, R.I., Lori, A., Randell, S.H., Oren, M., and Jetten, A.M. (2004). Critical role of p63 in the development of a normal esophageal and tracheobronchial epithelium. Am. J. Physiol. Physiol. 287, C171–C181.

Deblandre, G. a, Wettstein, D.A., Koyano-Nakagawa, N., and Kintner, C. (1999). A two-step mechanism generates the spacing pattern of the ciliated cells in the skin of Xenopus embryos. Development 126, 4715–4728.

Didon, L., Zwick, R.K., Chao, I.W., Walters, M.S., Wang, R., Hackett, N.R., and Crystal, R.G. (2013). RFX3 Modulation of FOXJ1 regulation of cilia genes in the human airway epithelium. Respir. Res. 14, 1.

Dobin, A., Davis, C.A., Schlesinger, F., Drenkow, J., Zaleski, C., Jha, S., Batut, P., Chaisson, M., and Gingeras, T.R. (2013). STAR: Ultrafast universal RNA-seq aligner. Bioinformatics 29, 15–21.

Dobzanski, A., Khalil, S.M., and Lane, A.P. (2018). Nasal polyp fibroblasts modulate epithelial characteristics via Wnt signaling. Int. Forum Allergy Rhinol. 8, 1412–1420.

Driskell, R.R. (2004). Wnt-responsive element controls Lef-1 promoter expression during submucosal gland morphogenesis. AJP Lung Cell. Mol. Physiol. 287, L752–L763.

Dubaissi, E., Rousseau, K., Lea, R., Soto, X., Nardeosingh, S., Schweickert, A., Amaya, E., Thornton, D.J., and Papalopulu, N. (2014). A secretory cell type develops alongside multiciliated cells, ionocytes and goblet cells, and provides a protective, anti-infective function in the frog embryonic mucociliary epidermis. Development 141, 1514–1525.

Ferrer-Vaquer, A., Piliszek, A., Tian, G., Aho, R.J., Dufort, D., and Hadjantonakis, A.K. (2010). A sensitive and bright single-cell resolution live imaging reporter of Wnt/-catenin signaling in the mouse. BMC Dev. Biol. 10.

Gomperts, B.N. (2004). Foxj1 regulates basal body anchoring to the cytoskeleton of ciliated pulmonary epithelial cells. J. Cell Sci. 117, 1329–1337.

Hackett, N.R., Shaykhiev, R., Walters, M.S., Wang, R., Zwick, R.K., Ferris, B., Witover, B., Salit, J., and Crystal, R.G. (2011). The human airway epithelial basal cell transcriptome. PLoS One 6.

Harland, R.M. (1991). In Situ Hybridization: an improved whole-mount method for Xenopus embryos. Methods Cell Biol. 36, 685–695.

Hashimoto, S., Chen, H., Que, J., Brockway, B.L., Drake, J.A., Snyder, J.C., Randell,S.H., and Stripp, B.R. (2012). β-Catenin–SOX2 signaling regulates the fate of developing airway epithelium. J. Cell Sci. 125, 932–942.

Hayes, J.M., Kim, S.K., Abitua, P.B., Park, T.J., Herrington, E.R., Kitayama, A., Grow, M.W., Ueno, N., and Wallingford, J.B. (2007). Identification of novel ciliogenesis factors using a new in vivo model for mucociliary epithelial development. Dev. Biol. 312, 115–130.

Heasman, J., Kofron, M., and Wylie, C. (2000). Δ the early Xenopus embryo: A novel antisense approach. Dev. Biol. 222, 124–134.

Heijink, I.H., De Bruin, H.G., Van Den Berge, M., Bennink, L.J.C., Brandenburg, S.M., Gosens, R., Van Oosterhout, A.J., and Postma, D.S. (2013). Role of aberrant WNT signalling in the airway epithelial response to cigarette smoke in chronic obstructive pulmonary disease. Thorax 68, 709–716.

Hirsch, N., Zimmerman, L.B., and Grainger, R.M. (2002). Xenopus, the next generation: X. tropicalis genetics and genomics. Dev. Dyn. 225, 422–433.

Hogan, B.L.M., Barkauskas, C.E., Chapman, H.A., Epstein, J.A., Jain, R., Hsia, C.C.W., Niklason, L., Calle, E., Le, A., Randell, S.H., et al. (2014). Repair and regeneration of the respiratory system: Complexity, plasticity, and mechanisms of lung stem cell function. Cell Stem Cell 15, 123–138.

Hou, Z., Wu, Q., Sun, X., Chen, H., Li, Y., Zhang, Y., Mori, M., Yang, Y., Que, J., and Jiang, M. (2019). Wnt/Fgf crosstalk is required for the specification of basal cells in the mouse trachea. Development 146, dev171496.

Huang, Y.L., and Niehrs, C. (2014). Polarized Wnt signaling regulates ectodermal cell fate in Xenopus. Dev. Cell 29, 250–257.

Kjolby, R.A.S., and Harland, R.M. (2017). Genome-wide identification of Wnt/β-catenin transcriptional targets during Xenopus gastrulation. Dev. Biol. 426, 165–175.

Kolm, P.J., and Sive, H.L. (1995). Efficient Hormone-Inducible Protein Function in Xenopus laevis. Dev. Biol. 171, 267–272.

Königshoff, M., Balsara, N., Pfaff, E.M., Kramer, M., Chrobak, I., Seeger, W., and Eickelberg, O. (2008). Functional Wnt signaling is increased in idiopathic pulmonary fibrosis. PLoS One 3, 1–12.

Liao, Y., Smyth, G.K., and Shi, W. (2014). FeatureCounts: An efficient general purpose program for assigning sequence reads to genomic features. Bioinformatics 30, 923–930.

Love, M.I., Huber, W., and Anders, S. (2014). Moderated estimation of fold change and dispersion for RNA-seq data with DESeq2. Genome Biol. 15, 550.

Lu, P., Barad, M., and Vize, P.D. (2001). Xenopus p63 expression in early ectoderm and neurectoderm. Mech. Dev. 102, 275–278.

Mall, M.A. (2008). Role of Cilia, Mucus, and Airway Surface Liquid in Mucociliary Dysfunction: Lessons from Mouse Models. J. Aerosol Med. Pulm. Drug Deliv. 21, 13–24.

Malleske, D.T., Hayes, D., Lallier, S.W., Hill, C.L., and Reynolds, S.D. (2018). Regulation of Human Airway Epithelial Tissue Stem Cell Differentiation by β P300, and CBP. Stem Cells 36, 1905–1916.

Mi, H., Muruganujan, A., Casagrande, J.T., and Thomas, P.D. (2013). Large-scale gene function analysis with the panther classification system. Nat. Protoc. 8, 1551–1566.

Mucenski, M.L., Nation, J.M., Thitoff, A.R., Besnard, V., Xu, Y., Wert, S.E., Harada, N., Taketo, M.M., Stahlman, M.T., and Whitsett, J.A. (2005). β differentiation of respiratory epithelial cells in vivo. Am. J. Physiol. Cell. Mol. Physiol. 289, L971–L979.

Niehrs, C. (2012). The complex world of WNT receptor signalling. Nat. Rev. Mol. Cell Biol. 13, 767–779.

Pongracz, J.E., and Stockley, R.A. (2006). Wnt signalling in lung development and diseases. Respir. Res. 7, 1–10.

Quigley, I.K., and Kintner, C. (2017). Rfx2 Stabilizes Foxj1 Binding at Chromatin Loops to Enable Multiciliated Cell Gene Expression. PLoS Genet. 13, 1–29.

Quigley, I.K., Stubbs, J.L., and Kintner, C. (2011). Specification of ion transport cells in the Xenopus larval skin. Development 138, 705–714.

Reynolds, S.D., Zemke, A.C., Giangreco, A., Brockway, B.L., Teisanu, R.M., Drake, J.A., Mariani, T., Di, P.Y.P., Taketo, M.M., and Stripp, B.R. (2008). Conditional Stabilization of β-Catenin Expands the Pool of Lung Stem Cells. Stem Cells 26, 1337–1346.

Rock, J.R., Onaitis, M.W., Rawlins, E.L., Lu, Y., Clark, C.P., Xue, Y., Randell, S.H., and Hogan, B.L.M. (2009). Basal cells as stem cells of the mouse trachea and human airway epithelium. Proc. Natl. Acad. Sci. 106, 12771–12775.

Rock, J.R., Randell, S.H., and Hogan, B.L.M. (2010). Airway basal stem cells: a perspective on their roles in epithelial homeostasis and remodeling. Dis. Model. Mech. 3, 545–556.

Rock, J.R., Gao, X., Xue, Y., Randell, S.H., Kong, Y.Y., and Hogan, B.L.M. (2011). Notch-dependent differentiation of adult airway basal stem cells. Cell Stem Cell 8, 639–648.

Ruptier, C., De Gaspéris, A., Ansieau, S., Granjon, A., Tanière, P., Lafosse, I., Shi, H., Petitjean, A., Taranchon-Clermont, E., Tribollet, V., et al. (2011). TP63 P2 promoter functional analysis identifies Δ-catenin as a key regulator of ΔNP63 expression. Oncogene 30, 4656–4665.

Schmid, A., Sailland, J., Novak, L., Baumlin, N., Fregien, N., and Salathe, M. (2017). Modulation of Wnt signaling is essential for the differentiation of ciliated epithelial cells in human airways. FEBS Lett. 591, 3493–3506.

Sive, H.L., Grainger, R.M., and Harland, R.M. (2000). Early Development of Xenopus laevis - A laboratory manual. Cold Spring Harb. Protoc. 5.

Soares, E., and Zhou, H. (2018). Master regulatory role of p63 in epidermal development and disease. Cell. Mol. Life Sci. 75, 1179–1190.

Stubbs, J.L., Davidson, L., Keller, R., and Kintner, C. (2006). Radial intercalation of ciliated cells during Xenopus skin development. Development 133, 2507–2515.

Stubbs, J.L., Oishi, I., Izpisúa Belmonte, J.C., and Kintner, C. (2008). The forkhead protein Foxj1 specifies node-like cilia in Xenopus and zebrafish embryos. Nat. Genet. 40, 1454–1460.

Stubbs, J.L., Vladar, E.K., Axelrod, J.D., and Kintner, C. (2012). Multicilin promotes centriole assembly and ciliogenesis during multiciliate cell differentiation. Nat. Cell Biol. 14, 140–147.

Sun, D.I., Tasca, A., Haas, M., Baltazar, G., Harland, R.M., Finkbeiner, W.E., and Walentek, P. (2019). Na + /H + exchangers are required for the development and function of vertebrate mucociliary Epithelia. Cells Tissues Organs 205, 279–292.

Tan, F.E., Vladar, E.K., Ma, L., Fuentealba, L.C., Hoh, R., Hernan Espinoza, F., Axelrod, J.D., Alvarez-Buylla, A., Stearns, T., Kintner, C., et al. (2013). Myb promotes centriole amplification and later steps of the multiciliogenesis program. J. Cell Sci. 126, e1–e1.

Tilley, A.E., Walters, M.S., Shaykhiev, R., and Crystal, R.G. (2014). Cilia Dysfunction in Lung Disease. Annu. Rev. Physiol. 77, 379–406.

Tran, H.T., Sekkali, B., Van Imschoot, G., Janssens, S., and Vleminckx, K. (2010). Wnt/ -catenin signaling is involved in the induction and maintenance of primitive hematopoiesis in the vertebrate embryo. Proc. Natl. Acad. Sci. 107, 16160–16165.

Vladar, E.K., and Brody, S.L. (2013). Analysis of ciliogenesis in primary culture mouse tracheal epithelial cells (Elsevier Inc.).

Walentek, P. (2018). Manipulating and Analyzing Cell Type Composition of the Xenopus Mucociliary Epidermis. In Kris Vleminckx (Ed.), Xenopus: Methods and Protocols, Methods in Molecular Biology, pp. 251–263.

Walentek, P., and Quigley, I.K. (2017). What we can learn from a tadpole about ciliopathies and airway diseases: Using systems biology in Xenopus to study cilia and mucociliary epithelia. Genesis 55, 1–13.

Walentek, P., Beyer, T., Thumberger, T., Schweickert, A., and Blum, M. (2012). ATP4a Is Required for Wnt-Dependent Foxj1 Expression and Leftward Flow in Xenopus Left-Right Development. Cell Rep. 1, 516–527.

Walentek, P., Bogusch, S., Thumberger, T., Vick, P., Dubaissi, E., Beyer, T., Blum, M., and Schweickert, A. (2014). A novel serotonin-secreting cell type regulates ciliary motility in the mucociliary epidermis of Xenopus tadpoles. Development 141, 1526–1533.

Walentek, P., Beyer, T., Hagenlocher, C., Müller, C., Feistel, K., Schweickert, A., Harland, R.M., and Blum, M. (2015). ATP4a is required for development and function of the Xenopus mucociliary epidermis - a potential model to study proton pump inhibitor-associated pneumonia. Dev. Biol. 408, 292–304.

Walters, M.S., Gomi, K., Ashbridge, B., Moore, M.A.S., Arbelaez, V., Heldrich, J., Ding, B. Sen, Rafii, S., Staudt, M.R., and Crystal, R.G. (2013). Generation of a human airway epithelium derived basal cell line with multipotent differentiation capacity. Respir. Res. 14, 26–30.

Warburton, D., El-Hashash, A., Carraro, G., Tiozzo, C., Sala, F., Rogers, O., Langhe, S. De, Kemp, P.J., Riccardi, D., Torday, J., et al. (2010). Lung Organogenesis. In Current Topics in Developmental Biology, pp. 73–158.

Warner, S.M.B., Hackett, T.L., Shaheen, F., Hallstrand, T.S., Kicic, A., Stick, S.M., and Knight, D.A. (2013). Transcription factor p63 regulates key genes and wound repair in human airway epithelial Basal cells. Am. J. Respir. Cell Mol. Biol. 49, 978–988.

Zuo, W., Zhang, T., Wu, D.Z.A., Guan, S.P., Liew, A.A., Yamamoto, Y., Wang, X., Lim, S.J., Vincent, M., Lessard, M., et al. (2015). P63+ Krt5+ distal airway stem cells are essential for lung regeneration. Nature 517, 616–620.

