## Supplementary figures and images for "ΔN-Tp63 mediates Wnt/β-catenin-induced inhibition of differentiation in basal stem cells of mucociliary epithelia"

### FigS1

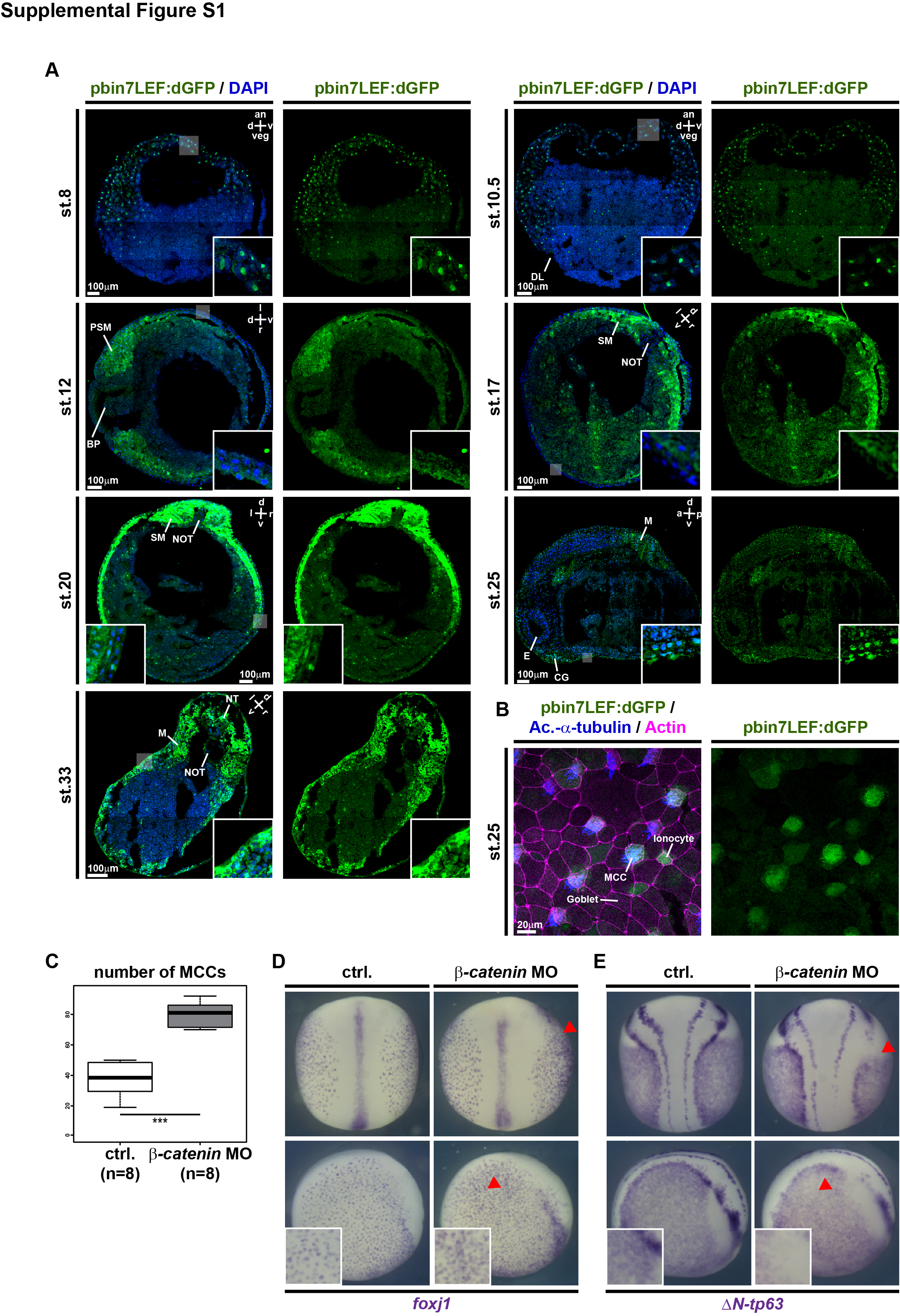

### FigS2

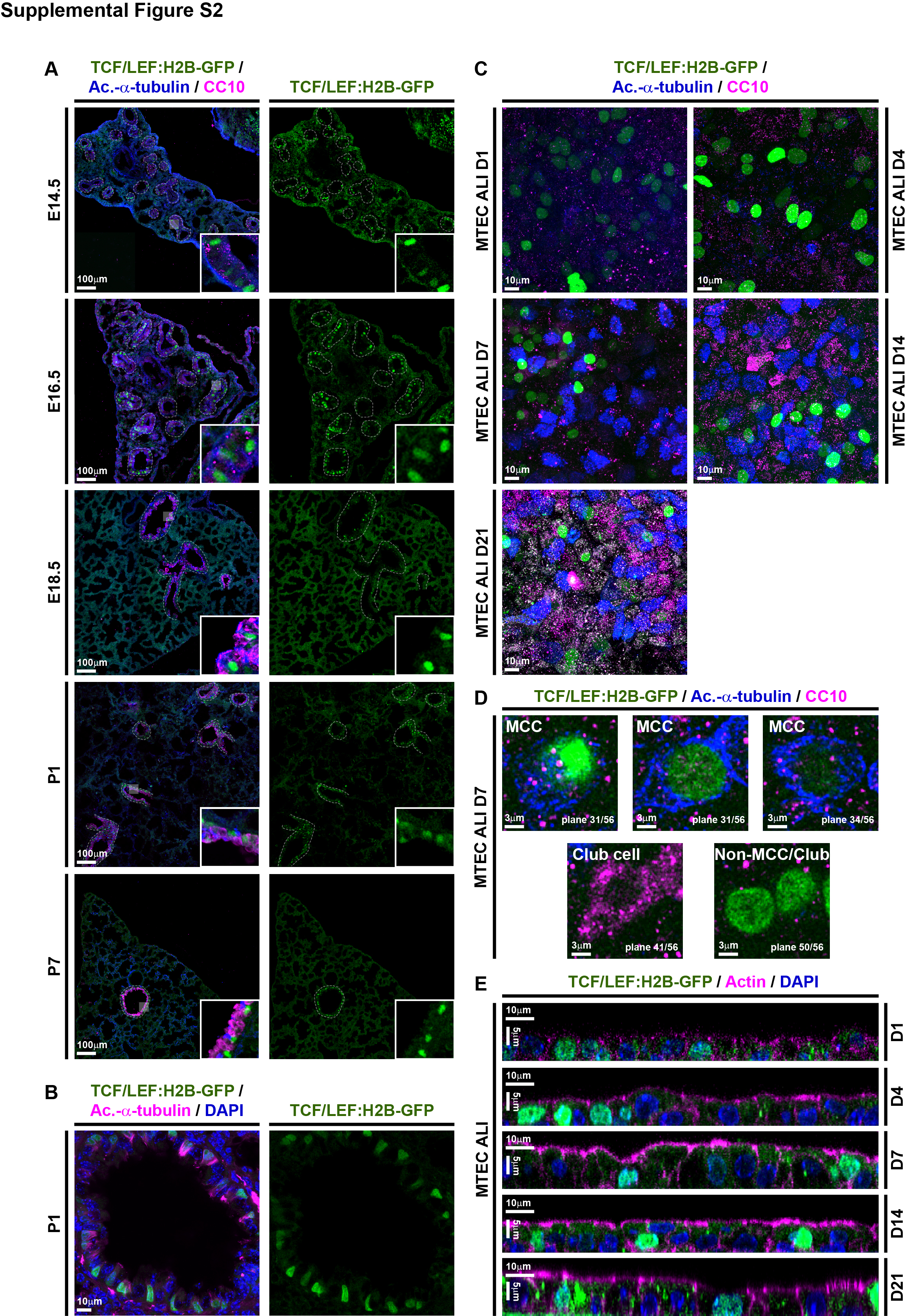

### FigS3

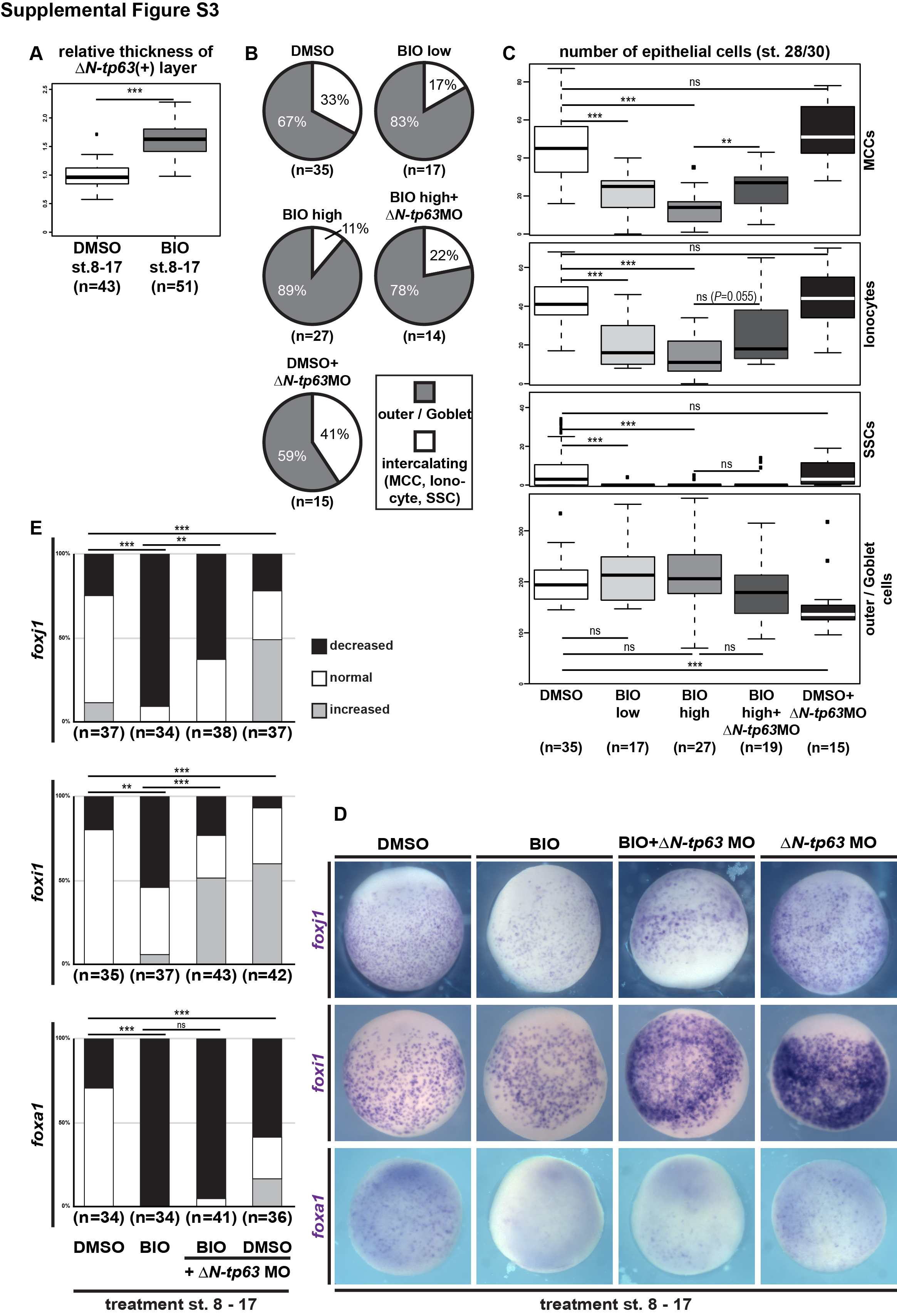

### FigS4

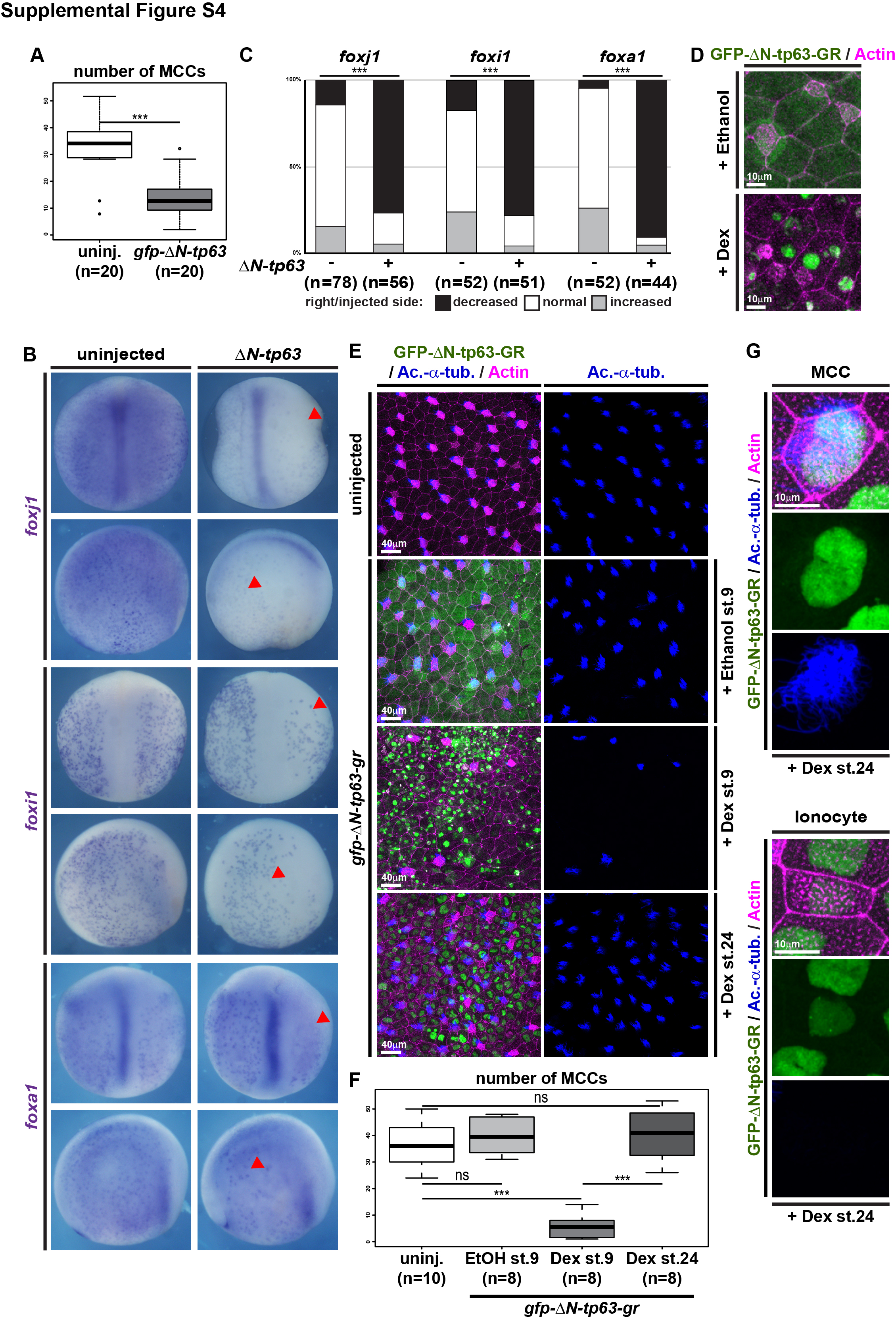

### FigS5

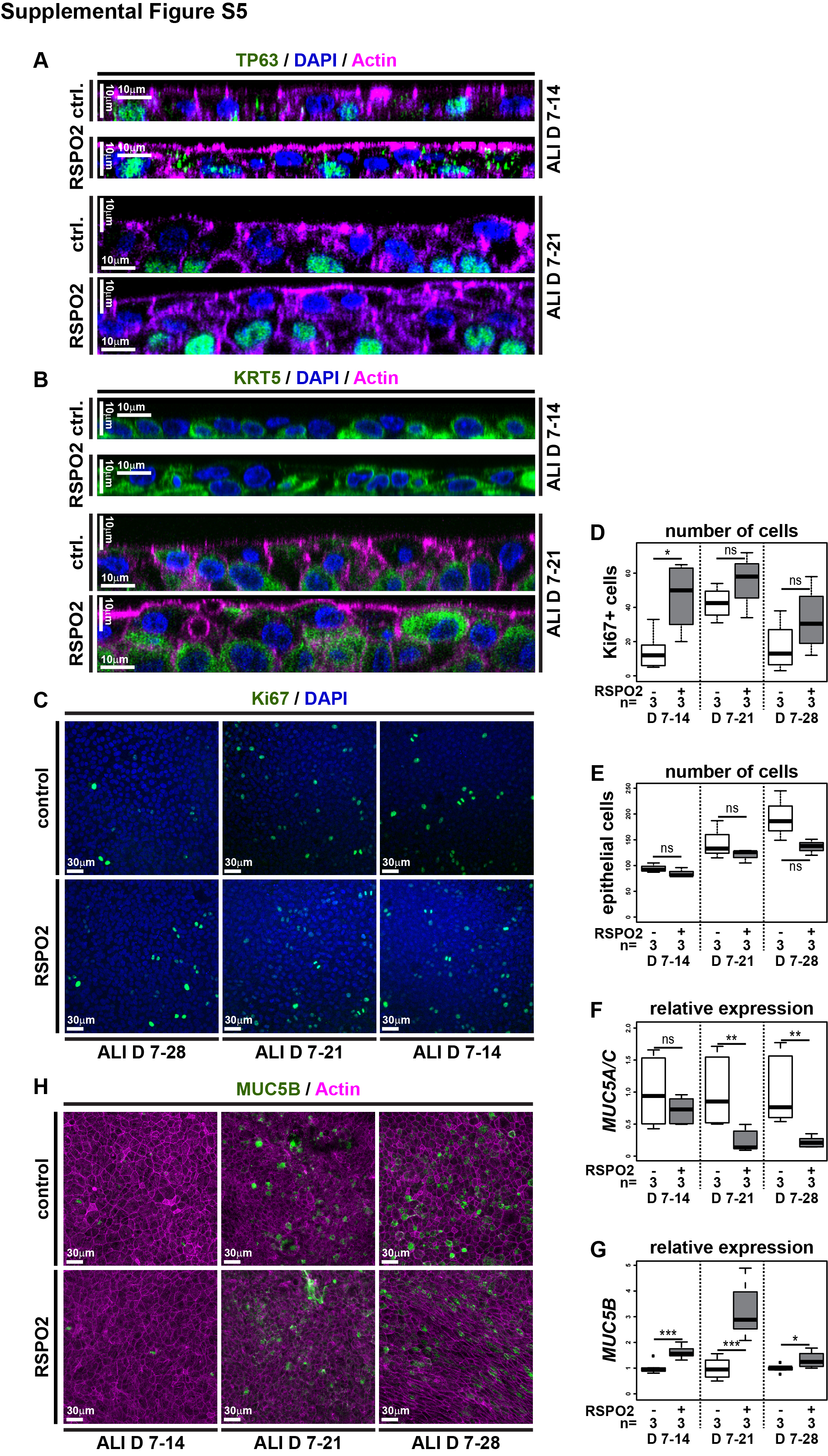

### FigS6

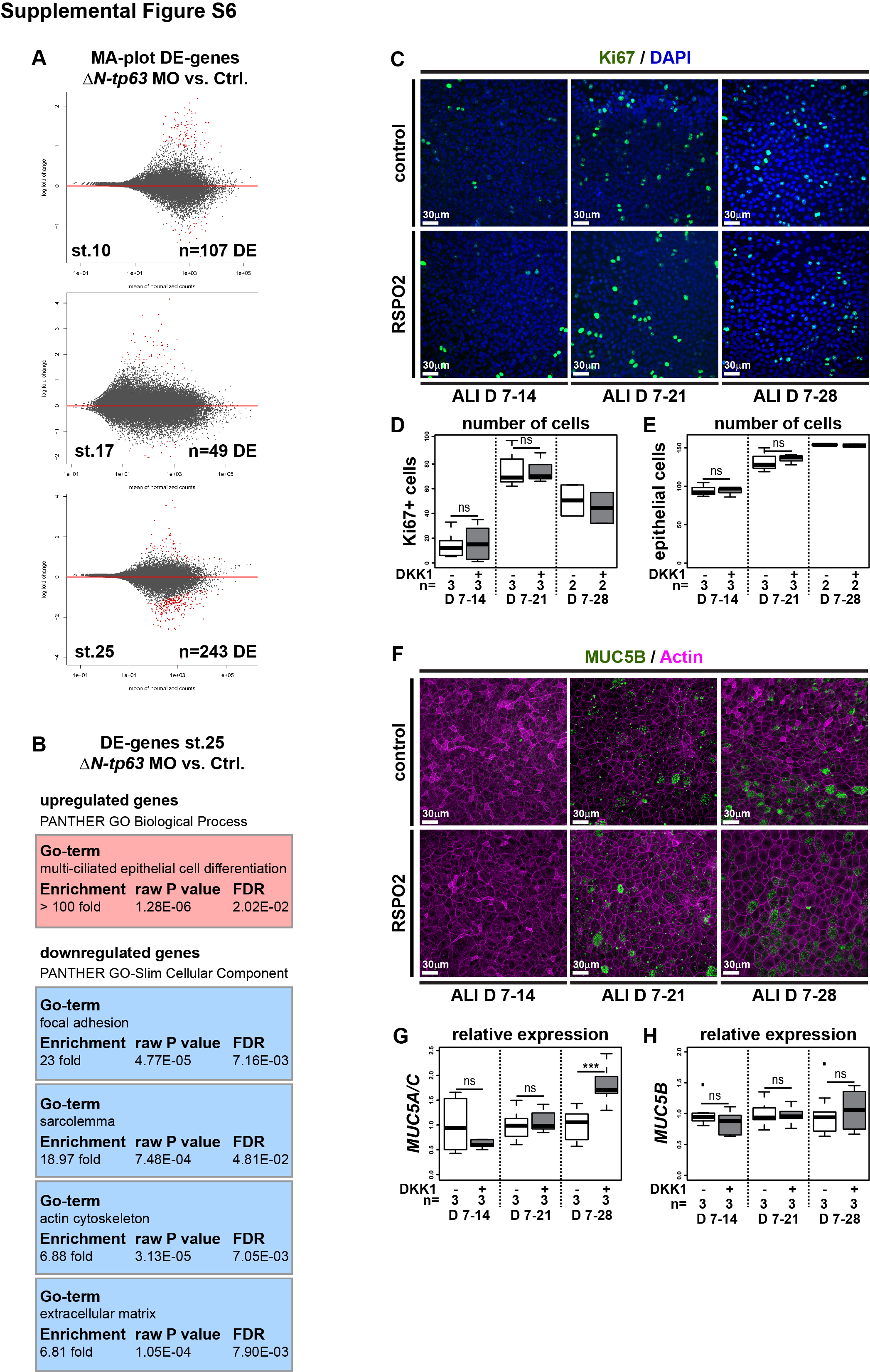

### FigS7

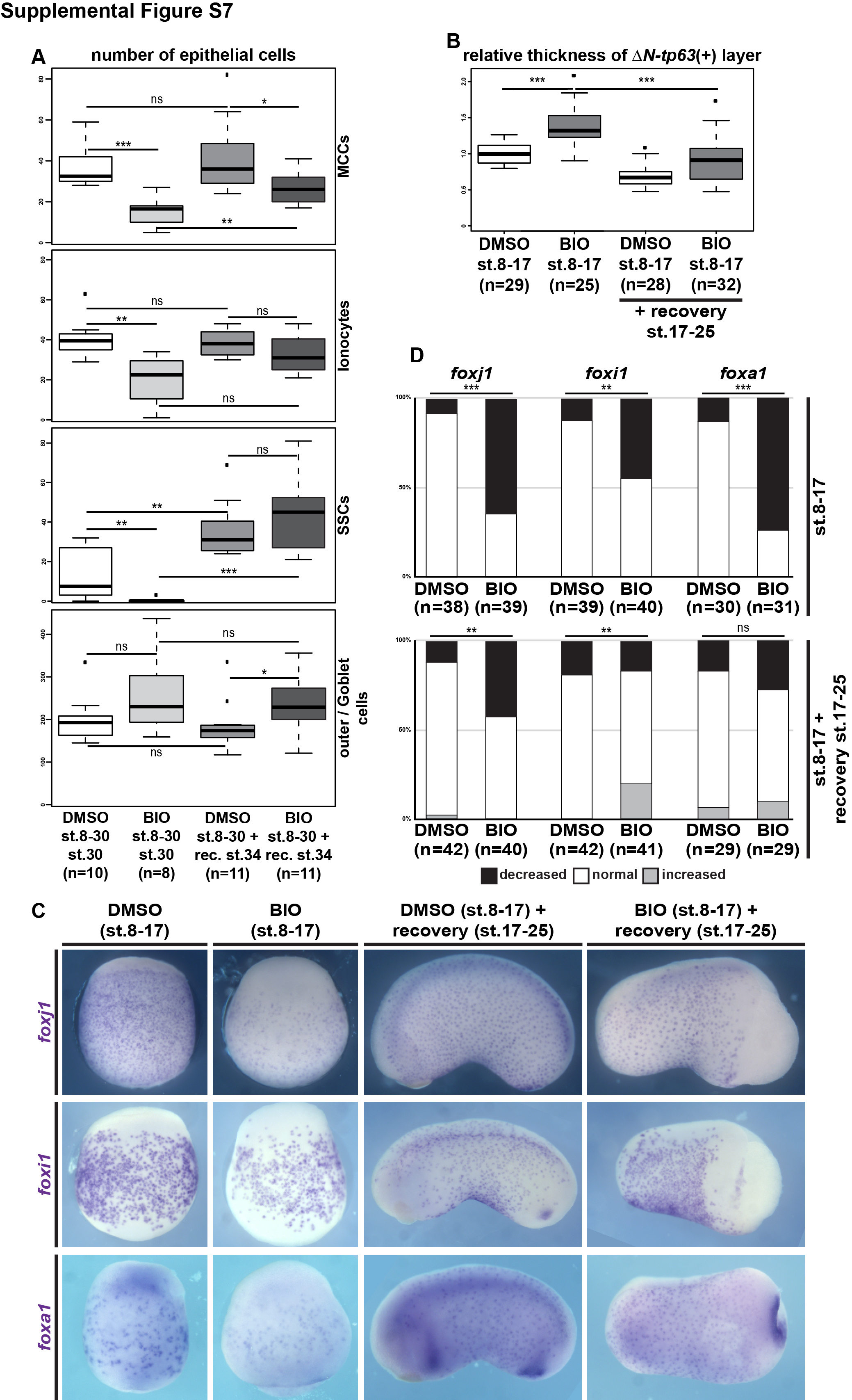

### FigS8

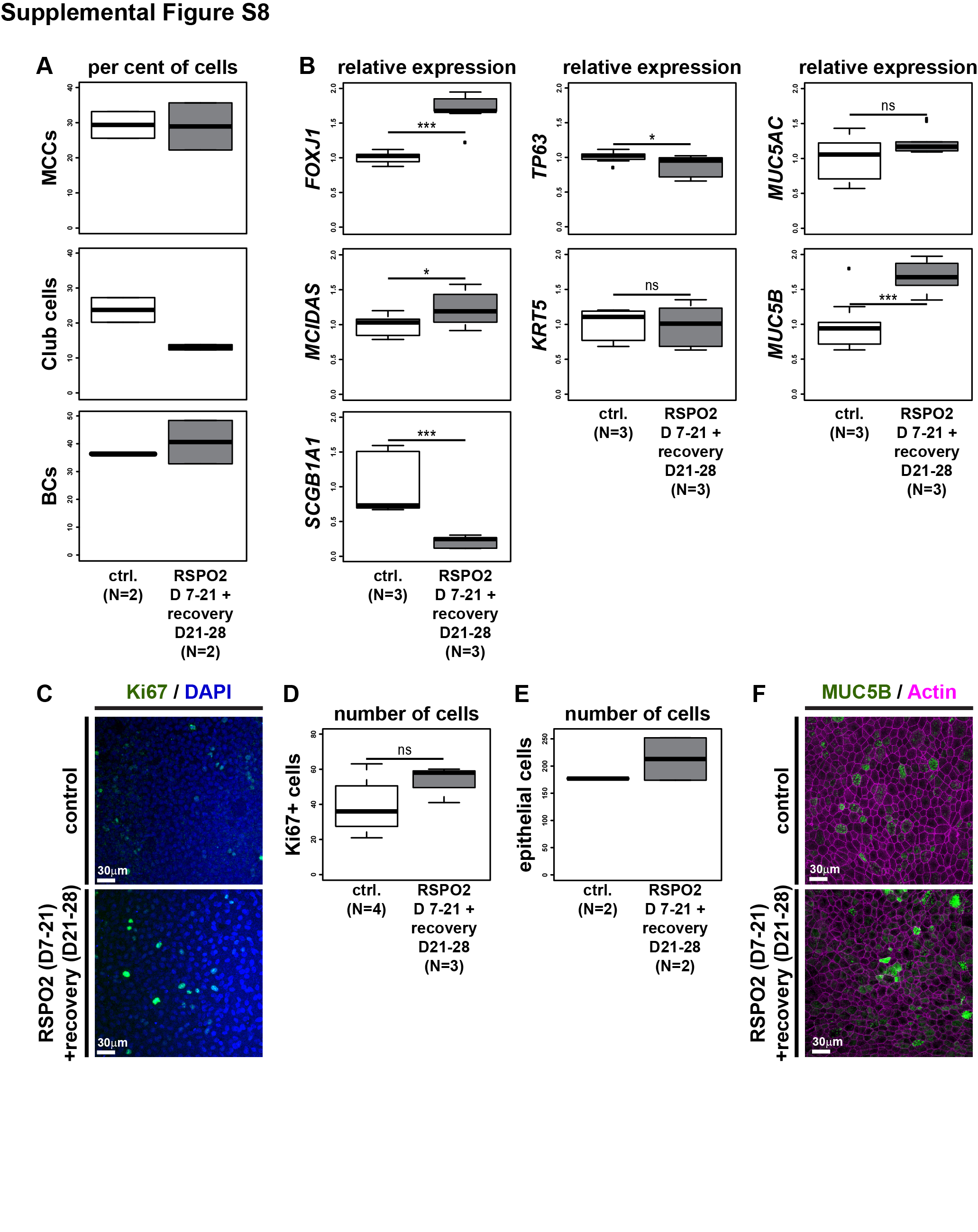
